# Heterozygous, polyploid, giant bacterium, *Achromatium*, possesses an identical functional inventory worldwide across drastically different ecosystems

**DOI:** 10.1101/2020.06.06.138032

**Authors:** Danny Ionescu, Luca Zoccarato, Artur Zaduryan, Sina Schorn, Mina Bižić, Solvig Pinnow, Heribert Cypionka, Hans-Peter Grossart

**Affiliations:** Leibniz Institute of Freshwater Ecology and Inland Fisheries, Neuglobsow, Germany; Berlin Brandenburg Institute of Biodiversity, Berlin, Germany; Department of Microbiology and Ecosystem Science, University of Vienna, Vienna, Austria; Institute for Chemistry and Biology of the Marine Environment, Oldenburg, Germany; Potsdam University, Potsdam, Germany

**Keywords:** Achromatium, Giant bacteria, Polyploidy, Geographical distribution of Achromatium, Eco-evolutionary advantage, Heterozygous bacteria

## Abstract

*Achromatium* is large, hyperpolyploid and the only known heterozygous bacterium. Single cells contain ca. 300 different chromosomes with allelic diversity typical of entire bacterial communities. Surveying all publicly available sediment sequence archives, we show *Achromatia* are common worldwide, spanning temperature, salinity, pH, and depth ranges normally resulting in bacterial speciation. Nevertheless, *Achromatia* display no ecotypic phylogenetic signal and contain a, globally identical, complete functional inventory. *Achromatia* cells from differing ecosystems (e.g. freshwater vs. saline) are, unexpectedly, equally functionally equipped but differ in gene expression patterns by transcribing only relevant genes. We suggest environmental adaptation occurs by increasing the copy number of relevant genes across the cell’s hundreds of chromosomes, without losing irrelevant ones, thus maintaining the ability to survive in any ecosystem type. The functional versatility of *Achromatium*, and its genomic features, reveal alternative genetic and evolutionary mechanisms, expanding our understanding of the role and evolution of polyploidy in bacteria while challenging the bacterial species concept and drivers of bacterial speciation.

## Introduction

Bacteria are typically well adapted to their environment (Bleuven and Landry 2016) with different levels of tolerance to changes in ambient conditions (Häusler et al. 2014; Saarinen et al. 2018). Adaption to novel (micro)environments includes changing, removing or incorporating new genes (Wiedenbeck and Cohan 2011; Hottes et al. 2013; Salcher et al. 2019). Such adjustments are then evolutionary stabilized in the population allowing those bacteria with increased fitness to proliferate (Tomatis et al. 2008; Milner et al. 2019).

Polyploid bacteria, defined as those harboring ten or more copies of their genomes, may be able to practice population-level experimental genomics within an individual cell, i.e. experimenting with genomic modifications on some chromosomes while maintaining pre-established functionality on others (Mendell et al. 2008; Oliverio and Katz 2014; Markov and Kaznacheev 2016). Eventually, gene conversion (i.e. asymmetric recombination) stabilizes one allele, likely the one that provides increased fitness (Ludt and Soppa 2019). The clonality of genomes in polyploid bacteria has been rarely discussed. While some results point towards minor differences (Mendell et al. 2008; Salman-Carvalho et al. 2016), some studies aiming at high-coverage assembly of polyploid bacteria suggest other bacteria harbor non-identical chromosomes (Winkel et al. 2016).

Genome comparison studies examining natural populations of single bacterial species revealed contrasting patterns regarding intra-population genomic variability. On the one hand, members of the *Roseobacter* clade, harboring multiple species, show little functional heterogeneity across different habitats likely due to acquisition of genes through lateral gene transfer (Newton et al. 2010). In contrast *Prochlorococcus*, the smallest but most abundant marine phototroph, forms populations consisting of hundreds of genomically different strains (Kashtan et al. 2014). Similarly, the most abundant oceanic heterotroph, *Candidatus* Pelagibacter sp. (SAR11) (Grote et al. 2012), and the abundant freshwater *Actinobacteria* of the AC-clade (Ghylin et al. 2014) form as well multiple genomic clades. Nevertheless, none of these population-wise heterogenous cells, have intracellular allelic divergence as documented in *Achromatia* (Ionescu et al. 2017).

*Achromatium* is a large sulfur-oxidizing bacterium that harbors calcium carbonate bodies in its periplasmatic space (Schorn et al. 2020). It is known mainly from freshwater sediments (Gray et al. 1999; Babenzien et al. 2015), but recently also from saline ones (Mansor et al. 2015; Salman et al. 2015), suggesting that it has a broad environmental distribution. In these sediments, *Achromatia*, occupy the upper few cm with most cells concentrated in the upper 2 cm at the oxic-anoxic interface (Gray et al. 1999). *All* known *Achromatium* cells are polyploid (Salman et al. 2015; Ionescu et al. 2017). Contradicting the definition of classical polyploidy, the multiple chromosomes of *Achromatia*, are not identical (Ionescu et al. 2017) making it the first and as yet only heterozygous bacterium (Ludt and Soppa 2019). Additionally, individual cells of *Achromatium* harbor genomic diversity characteristic of entire communities (Ionescu et al. 2017).

In this study, we aimed to explore the possible eco-evolutionary advantages of large heterozygosity in *Achromatia*. We used large-scale data mining, meta- and single cell genomics, and metatranscriptomics to highlight the ubiquitous presence of *Achromatia* in aquatic sediments without evident ecosystem-based genomic differentiation. Subsequently, we present the metabolic potential of the universal *Achromatium*.

## Results

Data recruitment of *Achromatium* small rRNA subunit (i.e. 16S rRNA gene) and functional gene sequences (see Material and Methods) from raw amplicon data as well as metagenomic and metatranscriptomic analyses of sediments worldwide, show *Achromatia* are ubiquitously present in ecosystems that differ drastically in their physico-chemical properties (Fig. 1). This includes freshwater lakes, rivers, coastal and deep marine sediments as well as several terrestrial environments for which water presence is not reported.

**Figure 1.**
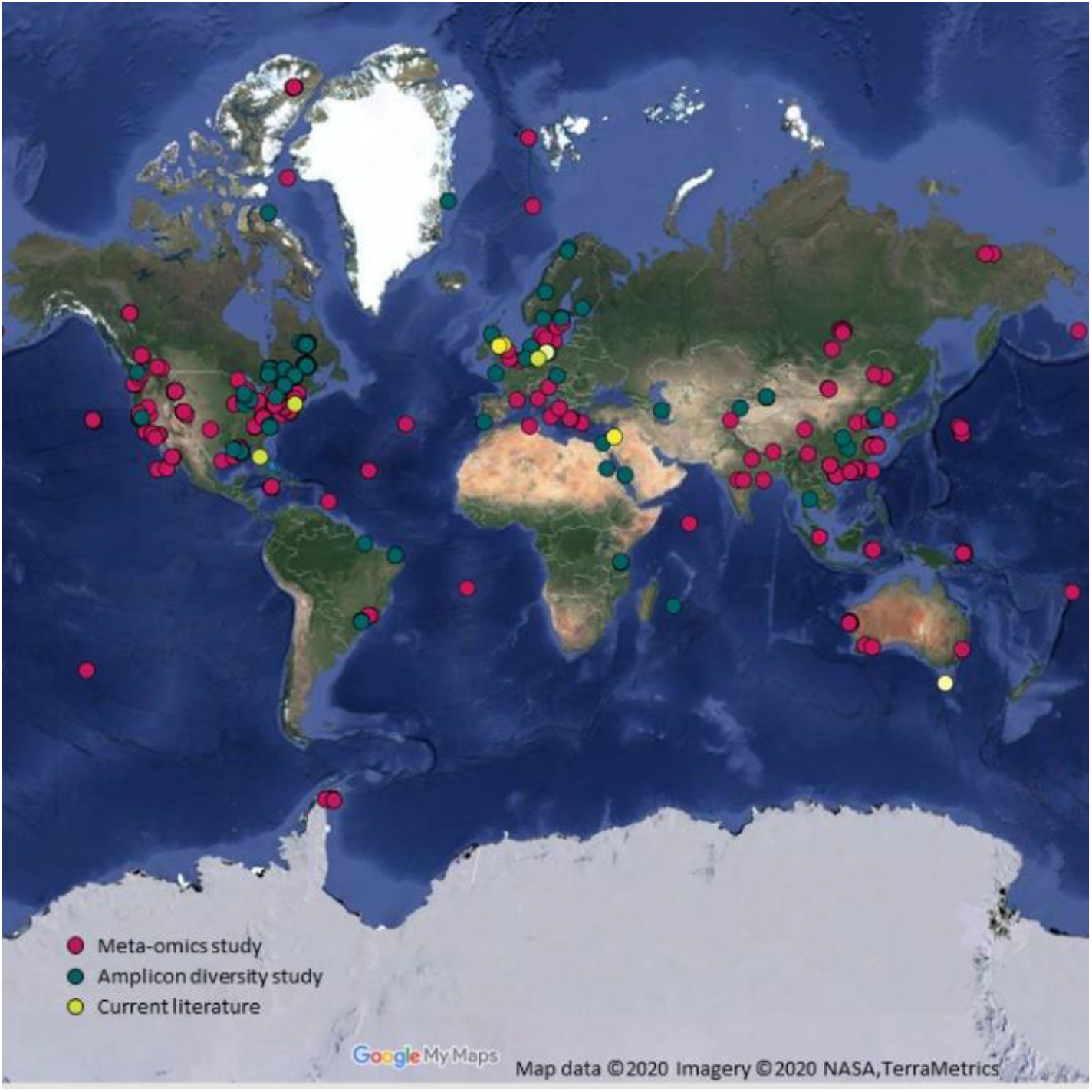
*Achromatia* are globally present in sediments of inland and oceanic waters as well as in extreme environments, e.g. hot springs, hypersaline lakes, the deep ocean, Arctic, and Antarctic ocean samples. The map was generated using sample metadata and the Google Maps website.

The upper and lower temperature, pH, salinity, and depth values for the ecosystems in which *Achromatia* were detected (including all sequence libraries surveyed in this study) are detailed in Table 1. These data highlight the ability of *Achromatia* to withstand a wide range of temperatures, pH values, salinity levels and hydrostatic pressures, though it is likely that in all these ecosystems it inhabits the sediment oxic-anoxic interphase niche.

**Table 1.** Lower and upper limits* of temperature, pH, salinity, and depth from which *Achromatia* sequences were recovered.

|  | Lower Limit | Upper Limit |
| --- | --- | --- |
| Temperature | 2 °C<sup>obs</sup> | 42 °C<sup>4</sup><br>60 °C<sup>2</sup> (101 °C) |
| pH | 4.7<sup>obs</sup> (4.2)<br>3.6<sup>1</sup> | 9.7<sup>1</sup> (10.3) |
| Salinity | Freshwater<sup>obs</sup> | 45 g L<sup>-1</sup><sup>5</sup> |
| Depth | 0 | 2600 m<sup>9</sup><br>4470 m<sup>1</sup> |

\*For studies which give a range of values without specifications on the exact sampling site, the more “extreme” value is given in parentheses. Numbers in uppercase show the number of different sequence libraries supporting these data with “obs” referring to direct observation of *Achromatia* cells.

We obtained 46,669 marine, 23,529 freshwater, 49 estuary and 89 terrestrial 16S rRNA gene sequences classified as *Achromatia*. As these originated from different amplicon or meta-omics studies that did not cover the same region of the gene, we chose the V4 region for further phylogenetic analysis as this was the one with the highest global coverage. To test whether, as expected, different habitats bare a phylogenetic signal, the sequences, dereplicated per ecosystem type (i.e. marine, freshwater, estuary, forest, soil or other), were used to construct a phylogenetic tree (Fig. 2, Fig. S1). Sequences of *Achromatia* do not exhibit any pattern, indicating the absence of any ecosystem-type based phylogenetic signal.

**Figure 2.**
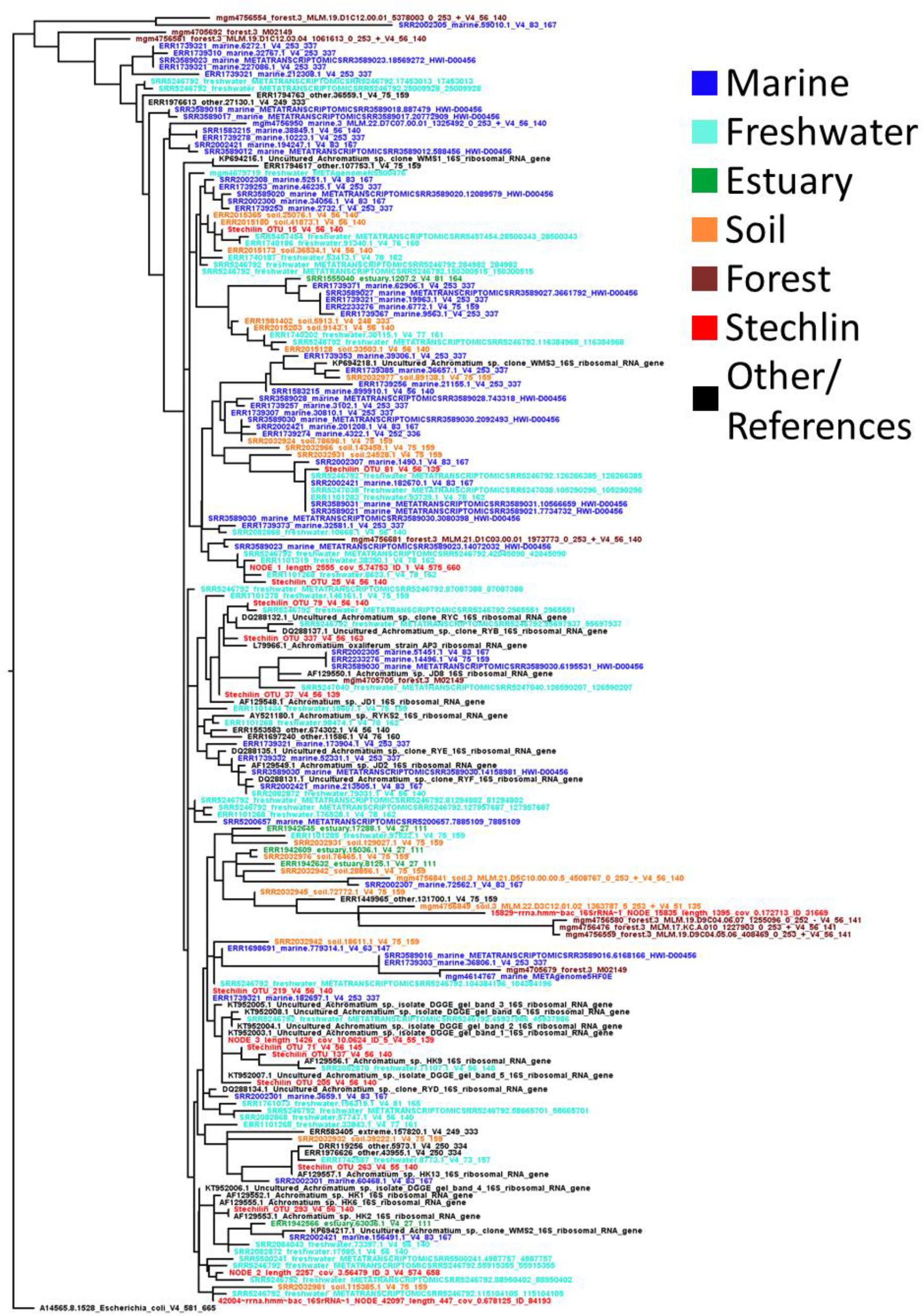
A phylogenetic tree constructed from the V4 region of 184 *Achromatia* 16S rRNA gene sequences recovered from raw amplicon, metagenome and -transcriptome sequence data deposited in various sequence archives, alongside reference sequences. To reduce the size of the tree 50 marine and 50 freshwater sequences were randomly chosen out of a larger dataset. The coloring highlights the lack of clustering based on ecological niches.

Though our analysis shows *Achromatia* in diverse ecosystems around the globe, the data recruited on *Achromatia* from extreme environments is scarce. However, the availability of many freshwater and marine sediment metagenomes and metatranscriptomes permits us to conduct a more in-depth examination of *Achromatia* on both sides of the salinity barrier.

Proteomic adaptation to salinity is typically reflected in the isoelectric point of proteins (Oren 2013). As such the distribution of calculated isoelectric points for proteins of a freshwater bacterium tends toward alkaline pH while of marine or halophilic organisms towards acidic pH (for a thorough comparison see Cabello-Yeves and Rodriguez-Valera, 2019). Such clear separations are obvious, for example, for the *Pelagibacteraceae* (Fig. 3A). Other phylogenetically closely related organisms (i.e. same genus) such as *Synechococcus* or members of the family *Beggiatoaceae* show less pronounced differences yet are generally more inclined towards acidic proteomes (Fig. S2). In contrast, our comparison of all known genomes of *Achromatia* from freshwater and saline (marine) ecosystems reveals neither major differences between the calculated acidity of their proteomes nor a strong inclination towards acidic proteomes (Fig. 3B). Though the proteomes of marine *Achromatia* have a lower percentage of basic proteins as compared to freshwater ones, the peak of the acidic isoelectric point of freshwater *Achromatia* is shifted towards a lower pH (Fig. 3B). Expanding the latter analysis to all *Achromatia* data recruited from freshwater, saline and intermediate (estuaries) ecosystems reflects the same phenomenon where no clear proteomic adaptation is observed with data from estuaries having a lower percentage of basic proteins (Fig. 3C). Interestingly, all *Achromatia* data shows a high number of proteins with a neutral isoelectric point (Fig. 3B-C).

**Figure 3.**
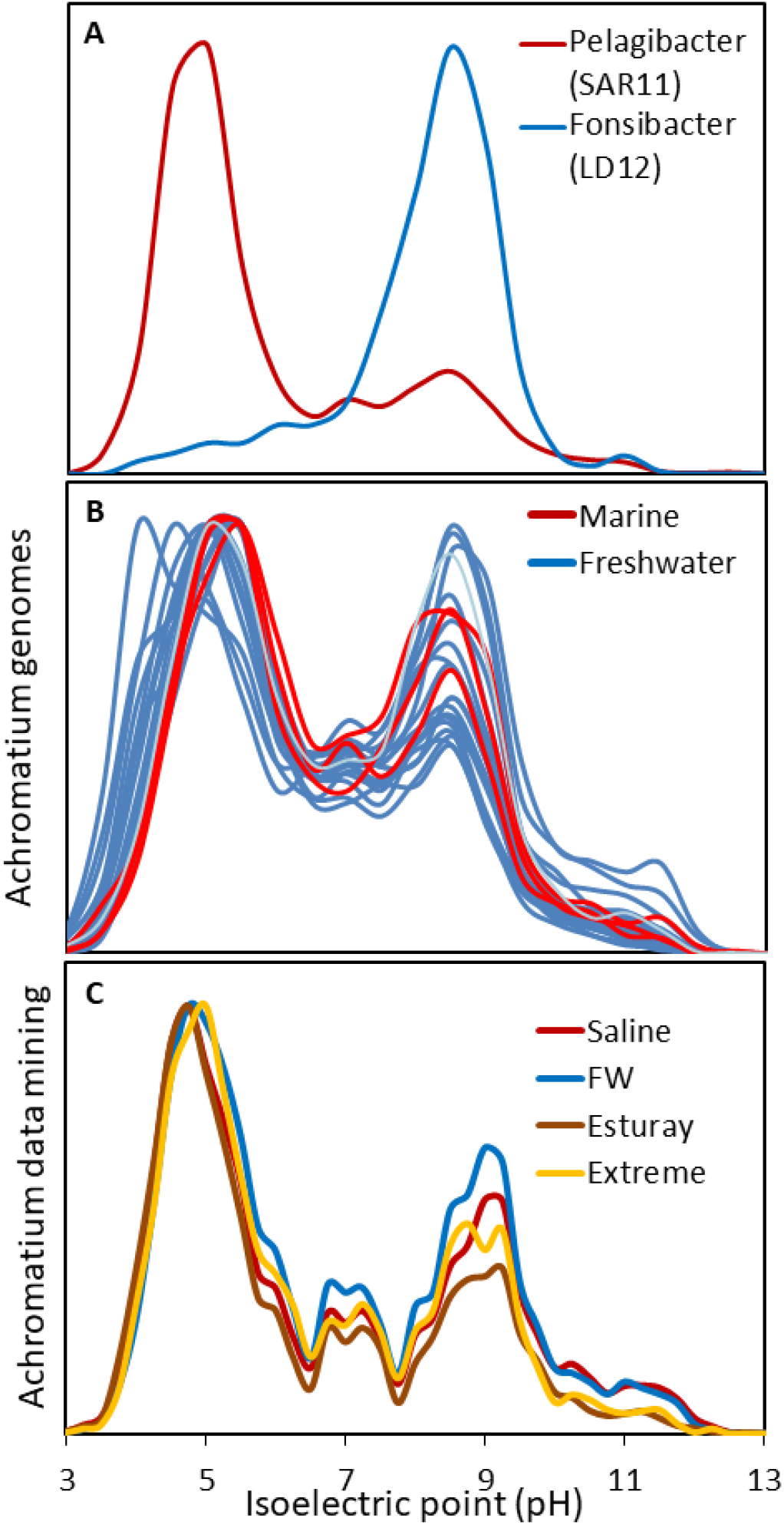
Isoelectric point histograms of two *Pelagibacteraceae* species from marine and freshwater ecosystems (A), the available freshwater and marine *Achromatia* genomes (B), and functionally recruited *Achromatia* data from metagenome and metatranscriptomic data (C).

Using the available genomic data from freshwater and marine *Achromatia*, we compared the metabolic functions of the two, typically evolutionary separated, ecosystems (Fig. S3, Supplementary dataset 1). This comparison suggests that genomes from each ecosystem harbor several unique functions though most functions overlap between freshwater and marine *Achromatia* genomes. Nevertheless, the available genomic information is very limited, with a strong bias towards freshwater from where numerous existing single-cell partial genomes and metagenomes (Ionescu et al. 2017) exist, although from one single environment. In contrast, data from the marine ecosystems consists of only 4 partial single cell genomes (Mansor et al. 2015; Salman et al. 2016). Therefore, we used the entire available genomic data to recruit *Achromatia* gene sequences from publicly available metagenomes and metatranscriptomes. Sequences for which a probability of 70 % or higher that the mapping to a known *Achromatia* sequence is correct were kept for further analysis. Given that average amino acid identity between *Achromatia* from the same and different environments is as low as 65 % and 55 %, respectively (Ionescu et al. 2017), a match probability of 70 % and higher is to be considered a high confidence match. Nevertheless, the per-gene average and median match probabilities of the high-quality matches (i.e. those above 70 %) was 95 % and 96 %, respectively (See Fig. S4 for data distribution). Functional genes of *Achromatia* were obtained from 704 sediment metagenomic studies. Ecosystem-type based comparison (saline *vs*. freshwater) of functions indicates that none of the functions is unique to either of the two types (Fig. 4). Clustering of the samples is a result of both coverage and environment with the former explaining ca. 25 % of the variability and the latter ca. 8 %, as shown by Permanova analysis (p=0.001 for both). Clustering of the genes (Clusters numbered 1 to 8) based on their presence/absence in the metagenomic samples (Supplementary dataset 2) did not reveal any functional differences between ecosystem-types and is therefore reported more in detail in the supplementary material.

**Figure 4.**
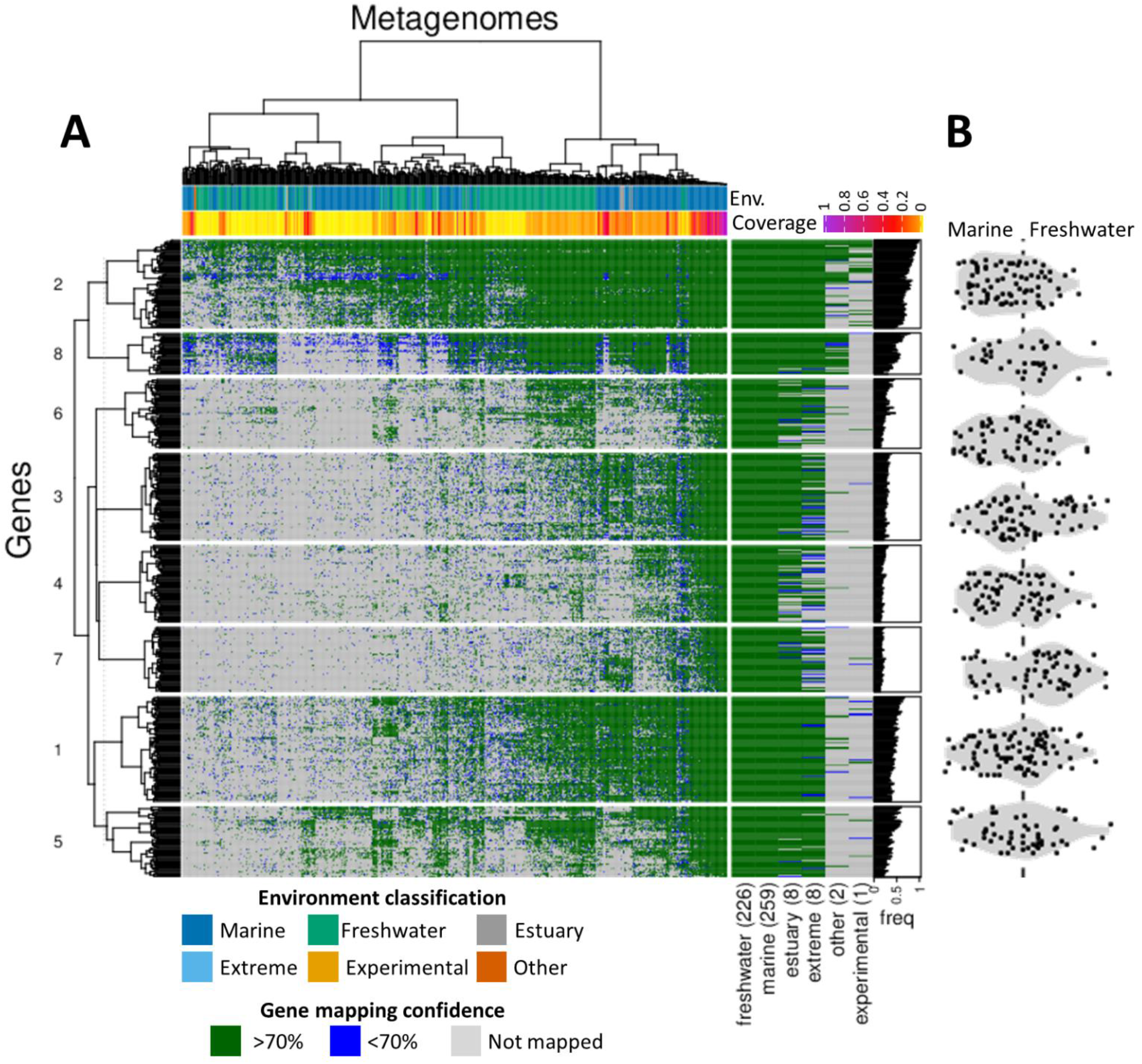
Functional potential of *Achromatia* from different sediment ecosystems as analyzed by mapping of raw sequence data to all known *Achromatia* annotated genes (A). When sequences from a study mapped positively to multiple alleles of an *Achromatium* gene, the mapping with the higher score was retained for presentation purposes. Green color shows a probability of 70 % or higher that the mapping to a known *Achromatia* sequence is correct. Blue shows a lower probability while gray indicates that the function was not detected in the sample. Given that average amino acid identity between *Achromatia* from the same and different environments is as low as 65 % and 55 %, respectively (Ionescu et al. 2017), a match probability of 70 % and higher is to be considered a high confidence match. Nevertheless, the per-gene average and median match probabilities of the high-quality matches (i.e. those above 70 %) was 95 % and 96 %, respectively (See Fig. S4 for data distribution). Functions were clustered based on presence absence in the samples revealing some genes are more common than others. Clustering information is given in Supplementary dataset 2. Functional profiles for overall freshwater and marine samples fully overlap with samples from estuaries, extreme environments (hot springs, hypersaline lakes, and soda lakes), and experiments containing less data. The latter are data resulting from bioreactor experiments carried out with marine sediment (see supplementary dataset 3). The per-sample sum of the fold-coverage for each known *Achromatia* protein, as calculated by the BBMAP mapping program, was used as a proxy for *Achromatia* sequence coverage in the sample. The ratio between a gene’s frequency in freshwater and marine ecosystems is presented for each cluster in a form of a violin boxplot (B). The 6-fold coverage differences between marine and freshwater (marine>freshwater) was accounted for in panel B by multiplying the freshwater gene presence frequency by the difference in median richness between the two ecosystem types (=1.3; see methods and Fig. S5).

To test whether *Achromatia* express genes differentially across various ecosystem types despite their globally identical functional potential (Fig. 4), we compared our transcriptomic data of *Achromatia* from Lake Stechlin to similar data recruited from public sediment metatranscriptomes (Fig. 5). Unlike for the metagenomic data, the samples, clustered according to gene coverage, generally group according to the different ecosystem types, separating freshwater from marine ones, with few exceptions. Nevertheless, similarly to the functional clustering of the metagenomic data, the gene clustering resulted in 11 groups that did not reveal any clear functional differences between *Achromatia* from the different ecosystem types. Clusters number 4 and 11 (Fig. 5), consisting mostly of genes involved in metabolism, signal processing and genetic information processing are expressed across both marine and freshwater ecosystems, though to a higher extent in freshwater. Cluster 6 is also expressed across all ecosystems. However, a higher expression is observed in two marine and a freshwater subset. This cluster contains the dissimilatory sulfite reductase and the adenylyl sulfate reductase, both typically involved in sulfur reduction and assimilation. Nevertheless, these enzymes are also found in sulfide oxidizers and function in reverse oxidizing intracellular sulfur, including in the closest known phylogenetic relative to *Achromatia, Allochromatium vinosum* (Dahl et al. 2005). Cluster 7 is expressed preferentially in the same subset of freshwater samples as cluster 6 (Fig. 5). It contains among others a sulfur transferase, another type of enzyme known to be involved in sulfur oxidation (Wang et al. 2019). Cluster 2 contains the largest number of genes most of which are involved in the central metabolism of the cell (Fig. 5). This cluster is mostly detected in our *Achromatia-targeted* metatranscriptomes and in a dataset of marine-sediment bioreactor experiments. It is likely that a higher expression of other genes combined with a relatively low sequencing depth of *Achromatia* in most studies led to the detection of these house-hold genes in these few high-coverage data sets.

**Figure 5.**
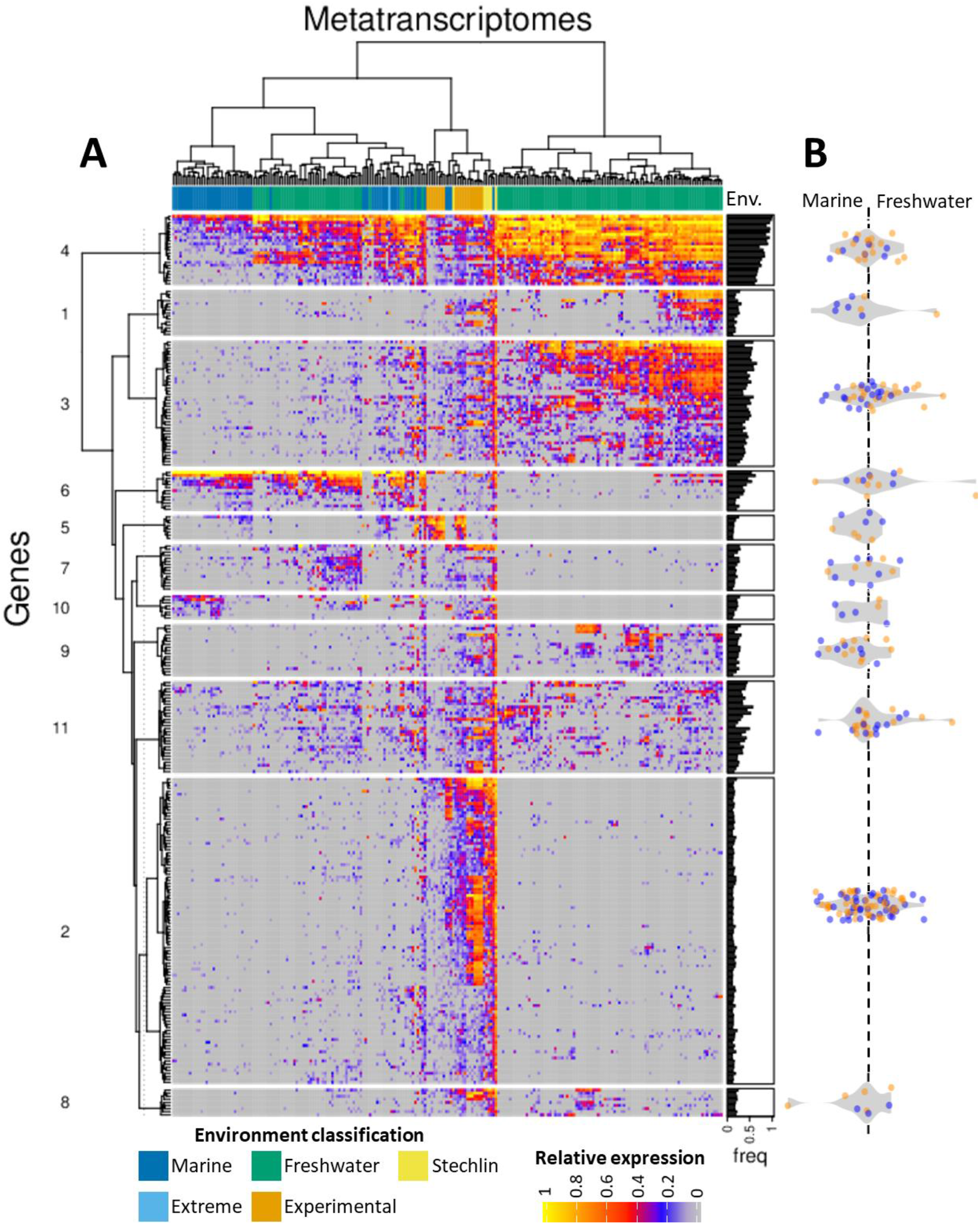
Analysis of ca. 300 publicly available sediment metatranscriptomes mapped to all known *Achromatia* functional genes and excluding ribosomal RNA and ribosomal proteins (A). To account for difference in sequencing depths and *Achromatia* cell abundance per sample, data was log-transformed and normalized per sample to range between 0-1. The ratio between a gene’s average non-zero expression in freshwater and marine ecosystems is presented for each cluster in a form of a violin boxplot (B) where symbols are colored according to the ecosystem type in which the gene was more frequently recovered in the metagenomes (i.e. Fig 4B). Thus, genes in orange and blue were more frequent in the freshwater and marine metagenomes, respectively. Zero values were omitted from the averaged expression to accommodate for coverage differences between studies.

Each gene in the transcriptome was analyzed for its preferential expression in freshwater or marine ecosystems (Fig. 5B). This analysis reveals two main features of *Achromatia*. First, while some genes appear to be preferentially expressed in one of the two ecosystem types, the shape and location of the violin boxplots across the marine/freshwater axis reveals that on average, most gene clusters are similarly expressed in both ecosystem types. Second, some genes found in higher frequency in either freshwater or marine ecosystems in the metagenomic data (Fig. 4) are preferentially expressed in the opposite ecosystem type. Both features, point to the globally uniform functional potential of *Achromatia*, suggesting, that preferential expression of genes is driven by local environmental factors (e.g. available electron donors and nutrients). The clustering of freshwater samples together with marine ones may be related to increased salinity in these waters due to drying out. Alternatively it may be driven by other factors which result in an overall similar expression pattern regardless of salinity.

*Achromatia* sequences were also found in thermal springs (Table 1). Therefore, as the GC content of bacteria was found not to be correlated with their optimal growth temperature (Hurst and Merchant 2001), we analyzed the known *Achromatia* genomes for proteomic adaptations typical to thermophilic bacteria. We focused on two main characteristics of thermophilic bacteria that distinguish them from mesophiles. First, a strong positive correlation between the relative abundance of Glutamate (Glu) and that of the pooled abundances of Lysine (Lys) and Arginine (Arg) (Tekaia et al. 2002). Second, a high ratio in the proteome between charged and polar amino acids (Kumar and Nussinov 2001; Suhre and Claverie 2003). To minimize the effect of incomplete protein assemblies we focused on proteins larger than 150 amino acids (340±202). Interestingly, the calculated *Achromatia* proteome has a low ratio of charged *vs*. polar amino acids but a significant correlation between the abundance of Glu and that of Lys+Arg (R=0.5) (Fig. S6). The correlation of the latter two improved the longer the analyzed proteins were, maximizing at R=0.76. When the amino acid frequencies of the *Achromatium* proteome were compared to those typical for aquatic and terrestrial bacteria (Fig. S6), no common pattern emerged. *Achromatium* showed a slightly higher abundance of Leucine (Leu), Proline (Pro), Methionine (Met) and Isoleucine (Ile) and a much lower abundance of Glycine (Gly), Glutamine (Gln), Glutamate (Glu), Serine (Ser), Asparagine (Asn) and Aspartate (Asp) than other bacteria. Additionally, the amino acid frequency pattern of *Achromatium* was also not indicative of nitrogen or carbon limitation when compared to the model suggested by Hellweger et al. (2018).

Using the available genomic data and the functional overlap between *Achromatia* from different ecosystem types, we propose a general functional model for the *Achromatium* cell, regardless of the ecosystem in which it lives (Fig. 6). The model, based on KEGG modules and pathways, shows potential metabolic functions and highlights whether these could be confirmed with metatranscriptome data from Lake Stechlin. A full metabolic analysis of *Achromatium*, however, is beyond the scope of this study. Therefore, here, we bring forth only a few selected functions. The entire list of detected functions by the different means of annotation is provided in Supplementary dataset 2 alongside information on confirmed expression and clustering data matching the metagenome (Fig 4, S7) and -transcriptome (Fig 5, S8) data. Over 25 % of the proteins were assigned to a function involved in genetic information processing, both in the overall genomic data but also in the metatranscriptome data from Lake Stechlin. *Achromatium* likely harbors 3 out of the known 6 pathways for carbon fixation, the Calvin cycle, reductive TCA and reverse Acetyl CoA with calculated KEGG module completion levels in the genomes of 100 %, 90 % and 43 % confirmed by completion in the transcriptomics of 81 %, 90 % and 28 %, respectively. *Achromatium* misses a key gene for the classical reductive TCA pathway, the ATP-citrate lyase, however, it harbors and expresses 2-oxoglutarate carboxylase, an alternative gene that allows the process to be carried out (Aoshima and Igarashi 2006). *Achromatium* also possesses the ability to obtain carbon heterotrophically as suggested by the presence of genes for sugar metabolism and degradation of fatty acid and amino acids. *Achromatium* can potentially fix N_2_ into NH_3_ and lacks known transporters for nitrate or nitrite. As expected from a sulfur oxidizing bacterium, *Achromatium* can oxidize H_2_S to sulfate via the sulfate quinone reductase genes and possesses the SoX gene to utilize thiosulfate. Yet, the thiosulfate binding protein of the sulfate transporter could not be identified in the genomic data. Additionally, *Achromatium* possesses and expresses the sulfite reductases *dsrA* and *dsrB* genes possibly used as well in sulfide oxidation (Dahl et al. 2005). Elemental sulfur is evidently an intermediate as sulfur globules are typically seen in the cells, but the genes involved in the formation of sulfur globules envelopes (Pattaragulwanit et al. 1998) could not be identified in any *Achromatia* genome (Schorn et al. 2020). As previously suggested, *Achromatium* harbors the V-type ATPase (Salman et al. 2016) alongside the F-type ATPase.

**Figure 6.**
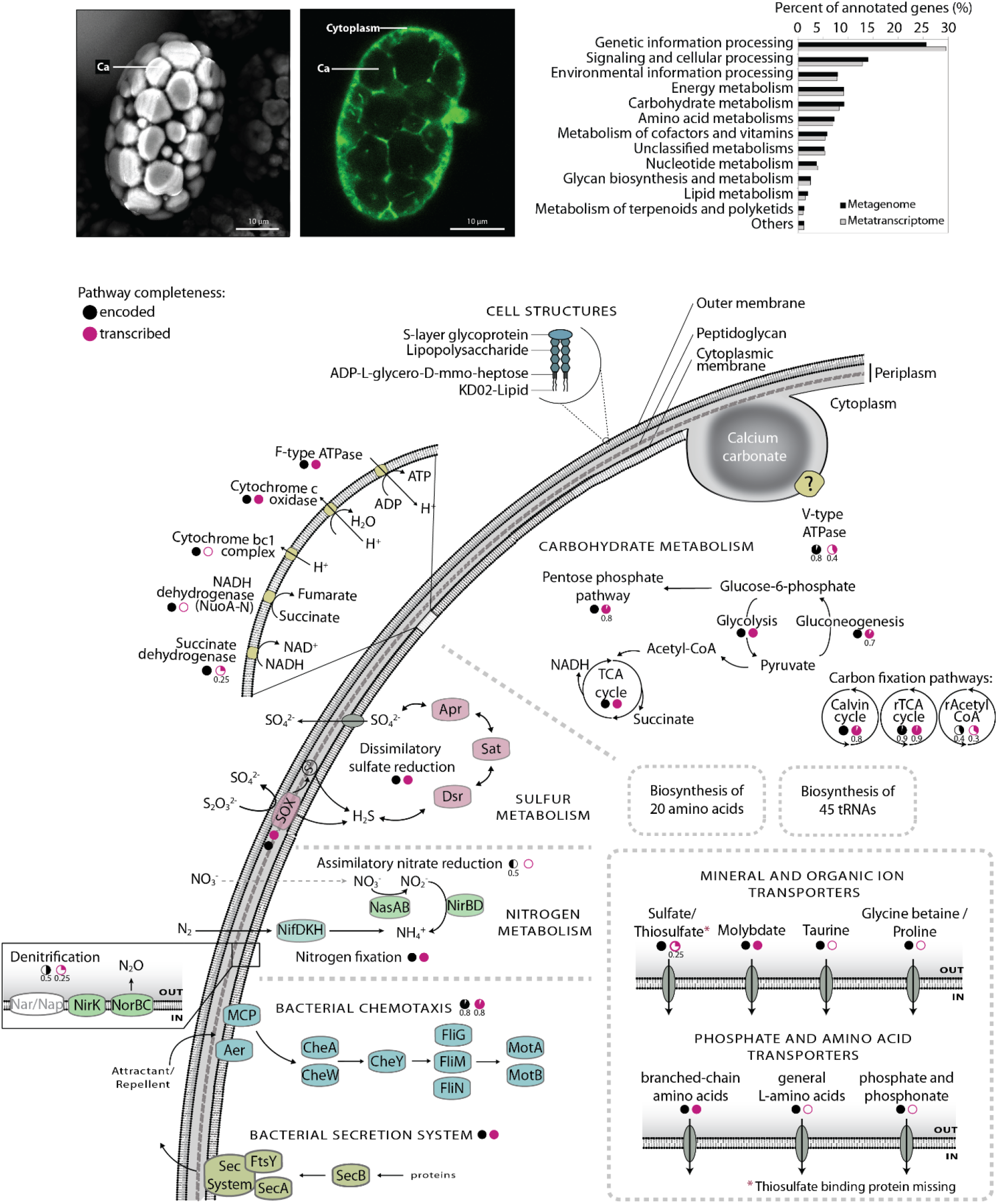
Main structural and functional characteristics of the universal *Achromatium* cell as deduced from KEGG modules and pathways as well as Pathway Tools analysis. The photomicrographs depict the general cell structure, where green color highlights the narrow cytoplasmatic space in between the Ca crystals. The fraction of genes annotated to different KEGG categories is shown as bar chart for both the genomic and transcriptomic data. Black and magenta pie-charts underneath pathways represent completeness in the published *Achromatia* genomes, and the transcriptomes obtained in this study, respectively. For incomplete pathways, the fraction present in the data is shown underneath the relevant circle. Pathways absent from the transcriptome are marked with an empty circle.

## Discussion

Most bacteria, in contrast to Eukaryotes, are known to streamline their genomes, minimizing over time the presence of genes that provide no advantage to their fitness (Lynch 2006; Koonin 2009; Bobay and Ochman 2017). Similarly, bacteria reduce the amount of non-coding DNA in their genome maintaining a relatively stable ratio between number of genes to non-coding regions and genome size to non-coding nucleotides (Giovannoni et al. 2005; Batut et al. 2014). Last, bacteria have developed effective mechanisms to avoid accumulation of deleterious mutations via Müller’s ratchet (Markov and Kaznacheev 2016) with gene conversion being the most common one (Ludt and Soppa 2019). Examples of genomic streamlining can be seen in genome reduction of symbiotic organisms (Boscaro et al. 2013), streamlining in oligotrophic environments (Giovannoni et al. 2005; Swan et al. 2013), or when transferring into such an environment (Salcher et al. 2019). Genomic adaptation can also be seen, for example, in homologous organisms when crossing the freshwater/salt barrier both in the genome (Zaremba-Niedzwiedzka et al. 2013; Tsementzi et al. 2019), the proteome (Cabello-Yeves et al. 2018) and metabolism (Walsh et al. 2013) as is the case of *Pelagibacterium* sp. (SAR11) and *Candidatus* Fonsibacter sp. (LD12) (Fig. 3A).

### *Achromatium*, a polyploid bacterium with ca. 300 chromosomes, challenges our understanding of general and genomic bacterial evolution

In a previous study (Ionescu et al. 2017), *Achromatia* were found to harbor an unprecedented degree of allelic divergence in a local population as well as in single cells covering all genes including those typically used as single cell markers. Recently, bacteria in the candidate phylum *Rokubacteria* (Becraft et al. 2017) have been found to harbor large genetic heterogeneity between single cells. However, whether like in the case of *Achromatia* these belong to a single species or they are functionally and phylogenetically diverging throughout this novel phylum, is not yet known. Ionescu et al., (2017) further found that unlike previously hypothesized for polyploid bacteria, the *Achromatia* chromosomes are not just replica of the same genome (Ionescu et al. 2017), rendering *Achromatia* the only known naturally heterozygous bacterium (Ludt and Soppa 2019). To explain this striking intracellular genomic variability, we previously proposed an evolutionary model of *Achromatia* (Fig. S9) that provided mechanistic hypotheses explaining the means by which such diversity can be generated (Ionescu et al. 2017). Two mechanisms were suggested to contribute to the generation of the large intracellular genetic diversity. The first, mobile genomic elements (transposons) which are abundant in *Achromatia* cells, contributes to genomic rearrangements resulting in inconsistent gene synteny and point mutations at insertion sites. The second, calcium carbonate cavities fill most of the cell volume, restricting the cytoplasmic volume to thin tubes or sheets between them (Schorn et al. 2020). This causes spatial separation of the chromosomes and the formation of genomic clusters which are possibly stabilized via gene conversion, as occurs in other polyploid prokaryotes (Soppa 2011; Ludt and Soppa 2019). However, due to the physical isolation of these clusters one from the other, different alleles may be fixed in each cluster. Upon cellular division, it was hypothesized that genomic clusters found in proximity to the division plane may be shuffled, with gene conversion resulting in the stabilization of new alleles and new gene synteny. This would result in two daughter cells that differ from each other, and from the mother cell. Both daughter cells still harbor genomic clusters that allow for continuous functionality alongside novel combinations enabling genetic experimentation, a trait that has been suggested as a benefit of bacterial polyploidy (Oliverio and Katz 2014; Markov and Kaznacheev 2016).

Here, we complement our theoretical reflections with large-scale data analyses to provide a better understanding of this unique eco-evolutionary phenomenon. Thus, we strive to understand the evolutionary changes *Achromatia* underwent to prevail in ecosystems characterized by phyisico-chemical parameters different enough to drive speciation rather than being occupied by a single species.

### *Achromatia* are abundant in aquatic sediments and are globally distributed

Earlier studies have already shown *Achromatia* to occur in contrasting freshwater ecosystems, e.g., oligotrophic Lake Stechlin and acidic bog lake Grosse Fuchskuhle (Glöckner et al. 1999), hinting towards a high metabolic versatility. Recently, it was also established that *Achromatia* are not present exclusively in freshwater but also in saline ecosystems (Mansor et al. 2015; Salman et al. 2015). These findings prompted us to conduct an extensive search for *Achromatium* 16S rRNA genes in all available amplicon, metagenome and -transcriptome-raw read libraries obtained from aquatic sediments. The results of this search, summarized in Fig. 1, demonstrate that *Achromatia* cells occur in almost all tested freshwater and marine samples including several unexpected ecosystem types such as hot-springs, hydrothermal vents, soda, and hypersaline lakes. Surprisingly, *Achromatia* were also found in samples of soils from non-aquatic ecosystems such as forests. Accordingly, *Achromatia* bridge a broad range of parameters selecting for specialized bacteria, such as temperatures favoring psychrophiles and thermophiles (<4 - >60 °C), alkaliphiles and acidophiles (pH <4 - >9.5), shallow-water organisms to piezophiles (0 - >3000 m depth) and marine and freshwater bacteria. These, extremely different, environmental parameters require special functional and genomic adaptations and accordingly drive speciation, i.e. the evolution of two or more species from a single common ancestor. Such is the case for example with freshwater, marine and thermophilic *Synechococcus* (Dvořák et al. 2014), freshwater and marine *Pelagibacteraceae, Candidatus* Pelagibacter sp. and *Candidatus* Fonsibacter sp. (Zaremba-Niedzwiedzka et al. 2013; Cabello-Yeves and Rodriguez-Valera 2019). Therefore, our findings rise the question whether various ecotypes can be distinguished genomically.

### *Achromatia* do not display an ecotypic phylogenetic signal

The presented data, mined from public read archives, were used to reconstruct the *Achromatia* phylogeny. Since most of the data originate from short read libraries, we used extracted sequences of the 16S rRNA gene variable regions for this purpose. A phylogenetic tree of the V4 region (Fig. 2), producing sequences from the largest number of different ecosystem types, clearly shows the lack of ecosystem-based phylogenetic separation. This is in contrast to what is known from other organisms which occur in different ecosystems such as marine and freshwater *Pelagibacteraceae* (Cabello-Yeves et al. 2018) or even in different ecological niches within the same ecosystem (Ahlgren and Rocap 2012). This is not entirely surprising given the fact that already in 22 single cells from one environment (Lake Stechlin, Germany), over 170 sequence variants were found (Ionescu et al. 2017). *Achromatia* stand out also from other large bacteria which occur in contrasting ecosystems and cluster phylogenetically accordingly (Salman et al. 2011; Teske and Salman 2014). These results suggest that if *Achromatia* harbor genetic adaptations to different ecosystem types, these are not reflected in the classic 16S rRNA phylogenetic marker.

### Freshwater *Achromatia* have proteomic adaptation to salinity

The freshwater/saline barrier is considered hard to cross with not many bacteria being able to move back and forth between the two types of aquatic systems (Walsh et al. 2013) for evolutionary significant periods of time (Bizic-Ionescu and Ionescu 2016). Bacteria inhabiting saline ecosystems evolved two main mechanisms to combat the higher osmotic pressure. The first, namely “Salt in” employs elevated intracellular concentration of ions (typically potassium instead of sodium) to match the external salt concentration. The second, “Salt out”, is more common at low to moderate salinities such as marine ecosystems. It employs elevated concentrations of small organic solutes such as glycine betaine, ectoine and trehalose, to account for the external salt concentration. The “Salt in” strategy necessitates an adapted proteome with an overall acidic isoelectric point (Oren 2013) which can function at high intracellular salt concentration. While the “Salt out” has no such requirements, most organisms inhabiting saline systems have adapted an acidic proteome or show an increased number of proteins with a low pH isoelectric point (Oren 2013; Cabello-Yeves and Rodriguez-Valera 2019).

We compared the predicted distribution of isoelectric points for all *Achromatia* proteins from genomes obtained from freshwater and saline systems and observed minimal difference between them (Fig. 2). A similar feature is observed when comparing the proteome assembled from the data mined from marine and freshwater sequence archives. In contrast, this analysis shows that both freshwater and marine *Achromatia* proteomes have a similar abundance of proteins with acidic and basic isoelectric points suggesting the cell maintain a permanent “readiness” for both freshwater and saline environments regardless of where they are. Interestingly, a similar pattern was observed in the proteome of other large bacteria from the family *Beggiatoaceae (Beggiatoa* spp. SS and PS) (Mußmann et al. 2007) obtained from the brackish water of the Baltic Sea (Fig. S2), and *Candidatus* Marithrix (Salman-Carvalho et al. 2016). The presence of osmolyte transport and synthesis systems in *Achromatia* cells from all ecosystem types, alongside sodium pumps and potassium channels further shows the cells are constantly prepared for higher or fluctuating salinities. As *Achromatia* cells were observed in sediments close to lake shores, it may be that these same adaptations help the cells survive increased salinities of drying freshwater environments.

### *Achromatium* is not an extremophile but may tolerate extreme conditions

Current literature places *Achromatia* in environments with moderate to low temperatures, neutrophilic to slightly basic environments and salinity up to that of seawater. In contrast, our survey shows *Achromatia* is present also in more extreme environments with temperatures exceeding 60 °C, pH reaching as low as 3.2 and salinity reaching 45 g L^-1^ (Table 1).

*Achromatia* have been mostly reported from neutral or alkaline environments with pH values above 7 (Head et al. 2000; Mansor et al. 2015), with one exception of the acidic Lake Grosse Fuchskuhle with a pH range (at the time of the study) of 4.2-4.7 (Glöckner et al. 1999). Our data survey shows the pH limits of *Achromatia* to be broader, ranging between 3.2 and 9.7. Multiple adaptations have been documented for bacteria enabling them to live in acidic environments (Mirete et al. 2017; Guan and Liu 2020) most of which revolve around structural adaptation of membranes to reduce proton permeability or mechanisms to increase proton export. As acid may damage macromolecules, some acid-tolerant microorganisms make additional use of chaperones (proteins that typically assist with the conformational folding or unfolding other proteins) to protect and repair such macromolecules. Of those the *Achromatia* pangenome contains the acid-stress protecting periplasmic chaperons DegP and SurA (Hong et al. 2012). Additionally, *Achromatia* harbor both V- and F-type ATPases which in case of acid stress can dissipate part of the proton gradient. Bacteria in acidic environments can also make use of buffering molecules such as lysine, histidine and arginine (Mirete et al. 2017). It was previously proposed (Yang et al. 2019) that the calcium carbonate crystals in the periplasmatic space of *Achromatia* may buffer the acidification effect of sulfide oxidation. In a similar manner, these crystals may provide a buffering mechanism in case of excess of protons in a low pH environment.

Elevated temperatures affect the bacterial membrane as well as the structure and stability of macromolecules such as proteins and nucleic acids. Accordingly, thermophiles, microorganisms with a preference to life at high temperatures, have acquired several adaptations resulting in improved thermostability of their membrane and protein structures (Kumar and Nussinov 2001). These changes come on top of efficient activity of chaperons and chaperonins (Richter et al. 2010). Nevertheless, thermotolerance, the ability to withstand temperatures higher than the organism’s typical environment can be acquired (Trent et al. 1994), *Achromatia* does not bare a clear thermophilic signature in the amino acid composition of its proteins. Therefore, it is likely that on a global scale it is not adapted to life at high temperatures with those populations found in thermal environments either relying on chaperone-based heat shock response or having locally adapted their proteomes. The latter, however, cannot be testes using the current dataset due to the limited number of samples recruited from extreme environments.

### *Achromatia* cells possess a full, identical, functional repertoire across different ecosystem types

Gene gain and loss upon transition between ecosystem types is a common phenomenon (Bleuven and Landry 2016; Mende et al. 2017; Milner et al. 2019; Salcher et al. 2019). Our global comparison of the functional inventory of *Achromatia*, shows that all functions identified so far in the published saline (Mansor et al. 2015; Salman et al. 2016) and freshwater (Ionescu et al. 2017) genomes are present in all ecosystem types (Fig. 4). A similar trend could already be observed when comparing the published genomes themselves (Fig. S2), however, several freshwater functions could not be detected in the marine genomes. This is likely an issue of coverage and data availability, given the genomic complexity of *Achromatium*, the sequencing depth necessary to properly assemble ca. 300, mostly different, chromosomes and the sole availability of 4 partial single-cell genomes from saline ecosystems. In contrast, data from freshwater systems includes a large metagenome, 6 single cells (Ionescu et al. 2017) and several more cells using Oxford NanoPore Technology (this study). Accordingly, the recovered unique functions (gene annotations) from the saline *Achromatia* genomes account individually for 17-41 % and combined for 53 % of the unique functions annotated from freshwater data. These numbers are however in par with the fractions obtained from individual bins (40±10 %) and single cells (42±3 %) of freshwater *Achromatia* from Lake Stechlin, highlighting the need for extensive sequencing depth to cover the full functional potential of *Achromatia* cells.

Gene detection frequency differs between saline, freshwater as well as other ecosystem types (Fig. 4B) despite the globally complete functional inventory of *Achromatia*. We attribute these differences to a mechanism allowing *Achromatia* to adapt to literally any environment. We hypothesize that, with time, genes more beneficial to the specific ecosystem type will occur in increasing numbers across the multiple chromosomes of *Achromatia* similarly to accumulation of beneficial gene duplications (Serres et al. 2009), while those of no immediate benefit will be “archived”. The extreme polyploidy of *Achromatia* cells and the hypothesized compartmentalization (Fig. S9; Ionescu et al., 2017), may allow *Achromatia* to conserve functions of no immediate benefit and at the same time maintain fitness in a given environment. This is in line with the already proposed advantage of bacterial polyploidy where these organisms can conduct “genetic experiments” while maintaining functionality (Mendell et al. 2008; Van de Peer et al. 2017). This is in addition to harboring extensive allelic divergence which likely allows *Achromatia* to fine tune their response to environmental changes, as observed in other organisms (Mock et al. 2017).

### *Achromatia* express genes out of their global pool based on the environment and immediate needs

Different expression patterns of *Achromatia* genes are observed in different ecosystem types, regardless of sequencing depth. This is not surprising, as organisms do not express all their genetic repertoire under all environmental conditions (Christie-Oleza et al. 2012). However, when comparing the averaged expression of functions in freshwater vs. those in saline systems, it becomes evident that functions detected preferentially in metagenomes from one ecosystem type can be more expressed in another. This further supports the presence of a global *Achromatia* function-inventory, from which required functions can be turned on if needed, despite having been, or *enroute* to being archived.

## Conclusions

We demonstrate a worldwide distribution of *Achromatia*, the only known heterozygous bacterium, across a broad range of ecosystem types. We show that *Achromatia* accumulate functions resulting in a globally identical functional pool. This is in addition to the previously documented allelic divergence. Accordingly, while even two daughter cells may differ in genomic sequence content and synteny - one from the other and from the mother cell, they harbor an identical functionality. We further propose that lowering the copy numbers of temporarily unnecessary genes in *Achromatia*, will not cause immediate functional loss. Hence, even if only a few copies of those genes will be maintained, the related functions can be recalled in case of need. The presence of *Achromatia* across a broad range of ecosystem types with drastically different characteristics suggests that the expectedly-high costs of generating, maintaining and regulating multiple, heterozygous full sets of genes regardless of the immediate environment, are paying off, providing *Achromatia* with the necessary adaptive power to survive anywhere. The, typical, localization of *Achromatia* in organic matter and nutrient rich upper cm of sediments, is likely a major factor in permitting such an expensive lifestyle. Similarly to the extreme allelic divergence (Ionescu et al. 2017), *Achromatium* challenges our understanding of genomic evolution in general and particularly that of polyploid organisms. In light of a plethora of large, polyploid bacteria (Ionescu and Bizic 2019), an urgent question that remains open is whether these, or other bacteria, share some or all of these newly discovered features.

## Material and Methods

### Collection of *Achromatia* cells

*Achromatia* cells were freshly collected from the shores of Lake Stechlin, NE Germany from the sediment surface at water depth of ca. 1 m. To obtain clean cells, collected sediments were sequentially passed through a series of mashes with pore sizes of 180 μm, 90 μm and 55 μm and collected in a large Petri dish. Subsequently, the filtrate was further cleaned under a binocular using a slow rotation movement which further separates *Achromatia* cells from fine sediments due to their different sedimentation properties. Sediment debris was removed and *Achromatia* cells were repeatedly cleaned using either RNA fixation buffer (see below) or lake water filtered through 0.22 μm diameter syringe filters. To remove epibionts, cleaned cells were washed for 30 min in 100 mM NaHCO_3_ buffer (Ionescu et al. 2017), a procedure which detaches the matrix of extracellular polymeric substances surrounding the *Achromatia* cells. These cells were further washed in buffer, sterile lake water and subsequently Milli-Q water prior to being directly processed for nucleic acid extraction. For single cell amplification, individual cells were collected with a thin glass pipette and sequentially cleaned and transferred using lake water filtered through a 0.1 μm pore size filter which was further autoclaved.

### RNA fixation

The need to manually enrich the *Achromatia* cells from the environment while preventing the degradation of cellular RNA or the expression of genes related to stress response, created several difficulties. Typical buffers as RNAlater, commercially available or lab-made using a saturated solution of ammonium sulfate, have low pH (<6) and a high salinity. The latter results in floating cells which cannot be separated from fine sediment using the sedimentation technique described above. Furthermore, the low pH leads to the rapid dissolution of the calcium carbonate bodies of *Achromatia* cells, leaving the cells invisible in the liquid. Consequently, based on the work by Lykidis et al., (2007) we used an modified Zinc-based buffer for the immediate fixation of cells. The final working recipe was as follows: 1% ZnCl_2_, 1% Zn(CF_3_COO)_2_, 0.05% Ca (CH_3_COO)_2_, 0.1 M Tris-HCl pH 8.0, 90 mM EDTA (Ethylenediaminetetraacetic acid), adjusted with NaOH to pH 8.0. The increased EDTA concentration prevented the salting out of the Zn at the high pH. Sediment collected for RNA worked was directly collected into the buffer, which was also used for all subsequent cleaning steps, except epibionts removal, as detailed above.

### RNA extraction

RNA was extracted from collected cells using a phenol/chloroform procedure adapted from (Nercessian et al. 2005) followed by DNA removal using the Turbo DNA fee kit (Thermo Fisher) as suggested in the manufacturer instructions. The cleaned RNA was stored in RNAstable tubes (Sigma) and sent for further processing to Mr. DNA (Molecular Research LP), Shallowater, Texas.

### RNA Sequencing

The RNA samples were resuspended in 25 μl of nuclease free water. RNA samples were cleaned using RNeasy PowerClean Pro Cleanup Kit (Qiagen). The concentration of RNA was determined using the Qubit^®^ RNA Assay Kit (Life Technologies). Whole transcriptome amplification was performed by using the QuantiTect Whole Transcriptome kit (Qiagen) followed by library preparation using KAPA HyperPlus Kits (Roche) following the manufacturer’s user guide. The concentration of double strand cDNA was evaluated (Table 1) using the Qubit^®^ dsDNA HS Assay Kit (Life Technologies). For six DNA samples, libraries were prepared using KAPA HyperPlus Kits (Roche). 25 ng DNA was used to prepare the libraries. Protocol starts with enzymatic fragmentation to produce dsDNA fragments followed by end repair and A-tailing to produce end-repaired, 5’-phosphorylated, 3’-dA-tailed dsDNA fragments. In adapter ligation step, dsDNA adapters are ligated to 3’-dA-tailed molecules. Final step is library amplification, which employs high fidelity, low-bias PCR to amplify library fragments carrying appropriate adapter sequences on both ends. Following the library preparation, the final concentration of all the libraries were measured using the Qubit^®^ dsDNA HS Assay Kit (Life Technologies), and the average library size was determined using the Agilent 2100 Bioanalyzer (Agilent Technologies). The libraries were then pooled in equimolar ratios of 2nM, and 8pM of the library pool was clustered using the cBot (Illumina) and sequenced paired end for 125 cycles using the HiSeq 2500 system (Illumina). The data is available at the Short Read Archive under project number PRJNA633541.

### DNA nanopore sequencing

Nanopore sequencing was conducted to improve the genome recovery of Achromatia from Lake Stechlin, as an addition to the data obtained in Ionescu et al (2017). To obtain the DNA concentrations necessary for NanoPore sequencing, single *Achromatium* cells were obtained as previously described (Ionescu et al. 2017) and amplified using the Repli-G kit (Qiagen, Hilden, Germany) following the manufacturer instructions. Libraries for NanoPore sequencing were then prepared using the LSK-108 kit following the manufacturer protocol, while excluding the size filtration. The prepared libraries were loaded on MIN107 R9 cells.

To incorporate the obtained reads into the previous assembly avoiding strand inversions incorporated by the phi29 polymerase in longer reads, the NanoPore reads were fragmented using the fastaslider.pl script from the Enveomics toolkit (Rodriguez-R and Konstantinidis 2016) into 250 nt long single end reads. These read were then co-assembled with the previous data using SPAdes assembler (Bankevich et al. 2012). However, no longer contigs than previously reported nor new functionality were obtained as a result from this assembly. The data is available at the Short Read Archive under project number PRJNA633773.

## Bioinformatic procedures

### Functional annotation

To generate a comprehensive functional database for *Achromatia* several tools were used. Prokka (Seemann 2014) was run with the most recent updates on all *Achromatia* available data from freshwater (Ionescu et al. 2017) and saline (Mansor et al. 2015; Salman et al. 2016) ecosystems. The generated GenBank files were subsequently used to generate metabolic models using Pathway Tools (v.23) (Karp et al. 2015; Karp et al. 2019). These models were generated for the different bins of *Achromatia* obtained from Lake Stechlin (Ionescu et al. 2017), for the genomes obtained from saline ecosystems (Mansor et al. 2015; Salman et al. 2016), as well as for a set of all unique *Achromatia* predicted genes.

The complete set of predicted *Achromatia* proteins was further annotated using the KEGG database using BlastKoala (Kanehisa et al. 2016) and KEGG module completeness was assessed using a custom script (available at https://github.com/lucaz88/R_script/blob/master/_KM_reconstruction.R).

Last, the complete set of *Achromatia* proteins was annotated using the dbCAN2 (Zhang et al. 2018) and CAZy (Cantarel et al. 2009) databases.

### RNAseq analysis

The RNAseq data was mapped against all available *Achromatia* genomic data to generate an *Achromatia* sequences pool, and subsequently against a pooled reference database consisting of all *Achromatia* annotated sequences from freshwater (Ionescu et al. 2017) and saline (Mansor et al. 2015; Salman et al. 2016) ecosystems using the BBmap tool (JGI, sourceforge.net/projects/bbmap/). Given *Achromatia* genomes contain multiple alleles for each function, for purpose of downstream analyses, the best mapping quality (mapq) per function was recorded from the resulting SAM files as well as the matching sequence coverage from the statistics file generated by the software.

### rRNA data mining and processing

To obtain novel *Achromatia* 16S rRNA sequences we first used the IMNGS online service (Lagkouvardos et al. 2016) to recruit data from raw-read amplicon libraries. Several runs of 10 query sequences (limit per run at the time) were submitted to the service using *Achromatia*, 16S rRNA available in the NCBI database as well as sequences generated in our previous work (Ionescu et al. 2017). Given the large variability between different *Achromatia* sequences (Ionescu et al. 2017), a similarity threshold of 90 % was set for the search. The raw amplicon library of the obtained results, were subsequently obtained and locally searched for *Achromatia* sequences using PhyloFlash (https://github.com/HRGV/phyloFlash) (Gruber-Vodicka et al. 2019). Sequences annotated as *Achromatia* were retained for further analysis. A list of the used studies is provided in supplementary dataset 2.

Additional 16S rRNA sequences were obtained from publicly available sediment metagenomes and metatranscriptomes deposited on the MG-RAST server or in the Short Read Archive (SRA). These data were downloaded and locally analyzed using PhyloFlash (https://github.com/HRGV/phyloFlash) (Gruber-Vodicka et al. 2019). Sequences annotated as *Achromatia* were retained for further analysis. A list of the used studies is provided in supplementary dataset 2.

The user-provided metadata was obtained from the repositories and used to classify the ecosystem type from where the sequence was obtained and to place the site on a global map in case relevant sequences were obtained. In cases where the ecosystem type was not properly reported, data was obtained from the matching publication if available or by using the sequence coordinates in Google Map.

Sequences were pooled according to their ecosystem type (saline, freshwater, extreme, river, estuary, other) and dereplicated. Since amplicon, metagenomic and metatranscriptomic sequences do not necessary overlap V-Xtractor (Hartmann et al. 2010) was used to extract the V1-V9 variable regions of the 16S rRNA. The variable regions sequences extracted from the environmental sequence pools were further dereplicated. Finally, random subsets of 50 sequences from freshwater and saline systems (mostly marine) were generated for phylogenetic tree reconstruction using the Fasttree 2 (Price et al. 2010) software with the GTR model and gamma correction.

### Metagenomic and metatranscriptomic public data processing

The downloaded data (MG-RAST and SRA, see above), was mapped against a pooled reference databased consisting of all *Achromatia* annotated sequences from freshwater (Ionescu et al. 2017) and saline (Mansor et al. 2015; Salman et al. 2016) environments using the BBmap tool (JGI, sourceforge.net/projects/bbmap/). Given *Achromatia* genomes contain multiple alleles for each function, for downstream analyses the best mapping quality (mapq) per function from the resulting SAM files, and the matching sequence coverage from the statistics file generated by the software, were recorded for each study.

The list of raw-read libraries used in this study and their available metadata in available as Supplementary dataset 3.

Heatmaps were plotted using the R packages ggplot2 (Wickham 2016) and ComplexHeatmap (Gu et al. 2016).

## Supporting information

Supplementary data set 2

Supplementary data set 3

Supplementary data set 1

## Acknowledgements

This work was supported by the German Federal Ministry of Education and Research (BMBF) through the BIBS (Bridging in Biodiversity Science; funding no. 01LC1501G) project. MB was funded through DFG project BI 1987/2-1. We thank Mr. Justin Stranz for his assistance.

## Supplementary Material

### Gene clustering from global metagenomic data

The cluster analysis highlights a cluster of *Achromatia* genes that are more commonly recovered from the data regardless of overall coverage and are hence either more abundant or less divergent than others (Fig. 4). Clusters 2 and 8 are the most ubiquitous ones and occur generally in samples from all environment types, followed by clusters 1 and 5. Cluster 2 and 1 are more diverse and consist of genes involved in carbohydrate, energy, nucleotide and amino acid metabolism alongside with genes involved in genetic information processing and a few tRNA genes. In contrast, cluster 8 contains almost exclusively a set of 33 tRNA genes. Cluster 5, while present in the four main environments (i.e. freshwater, marine, estuarine, and extreme habitats), is not common to all samples in those environments. This cluster contains additional genes involved in the central metabolism, but it additionally harbors several restriction enzymes and transposases. The latter have been previously shown to be abundant and diverse in freshwater *Achromatia* genomes (Ionescu et al. 2017) and likely play a significant role in the generation of the large intracellular allelic divergence observed in the *Achromatia* genome.

Cluster 6 is common only to a group of samples including a large subset of freshwater metagenomes. Among all genes in this cluster the potassium-transporting ATPase and the type I, multi-polypeptide enzymes NADH dehydrogenase (NADH-quinone oxidoreductase) stand out. The latter is differing from its single-polypeptide enzyme counterpart (Type II) by not having an iron-sulfur cluster.

Clusters 3, 4 and 7 are more typical in marine environments. However, as is the case with the other clusters, all genes are found in metagenomes from both marine and freshwater environments. This cluster harbors the genes for 2-oxoglutarate carboxylase, an alternative enzyme to ATP citrate lyase in the reductive TCA cycle (Aoshima and Igarashi 2006). This cluster further contains phosphate transporter genes (Pst), sodium pumps, and sensory genes involved in chemotaxis as well as additional, different, transposases and restriction enzymes.

**Figure S1.**
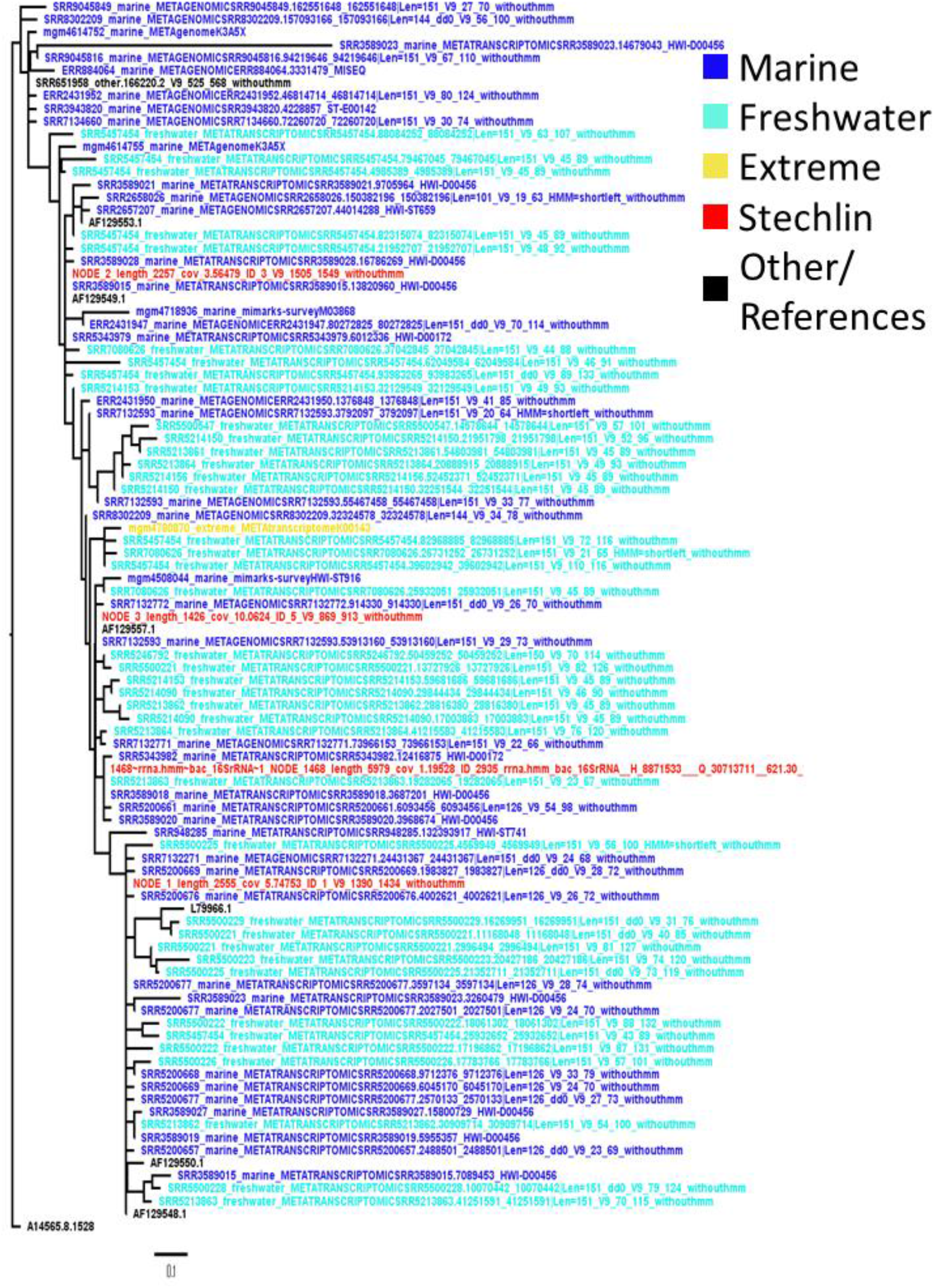
A phylogenetic tree constructed from the V9 region of *Achromatia* 16S rRNA sequences recovered from raw amplicon, metagenome and metatranscriptome sequence data deposited in sequence archives, alongside with reference sequences. Dereplication of the marine sequences reduced the set to 50 sequences and therefore 50 freshwater sequences were randomly chosen out of a larger dataset. The coloring highlights the lack of clustering based on ecological niches.

**Figure S2.**
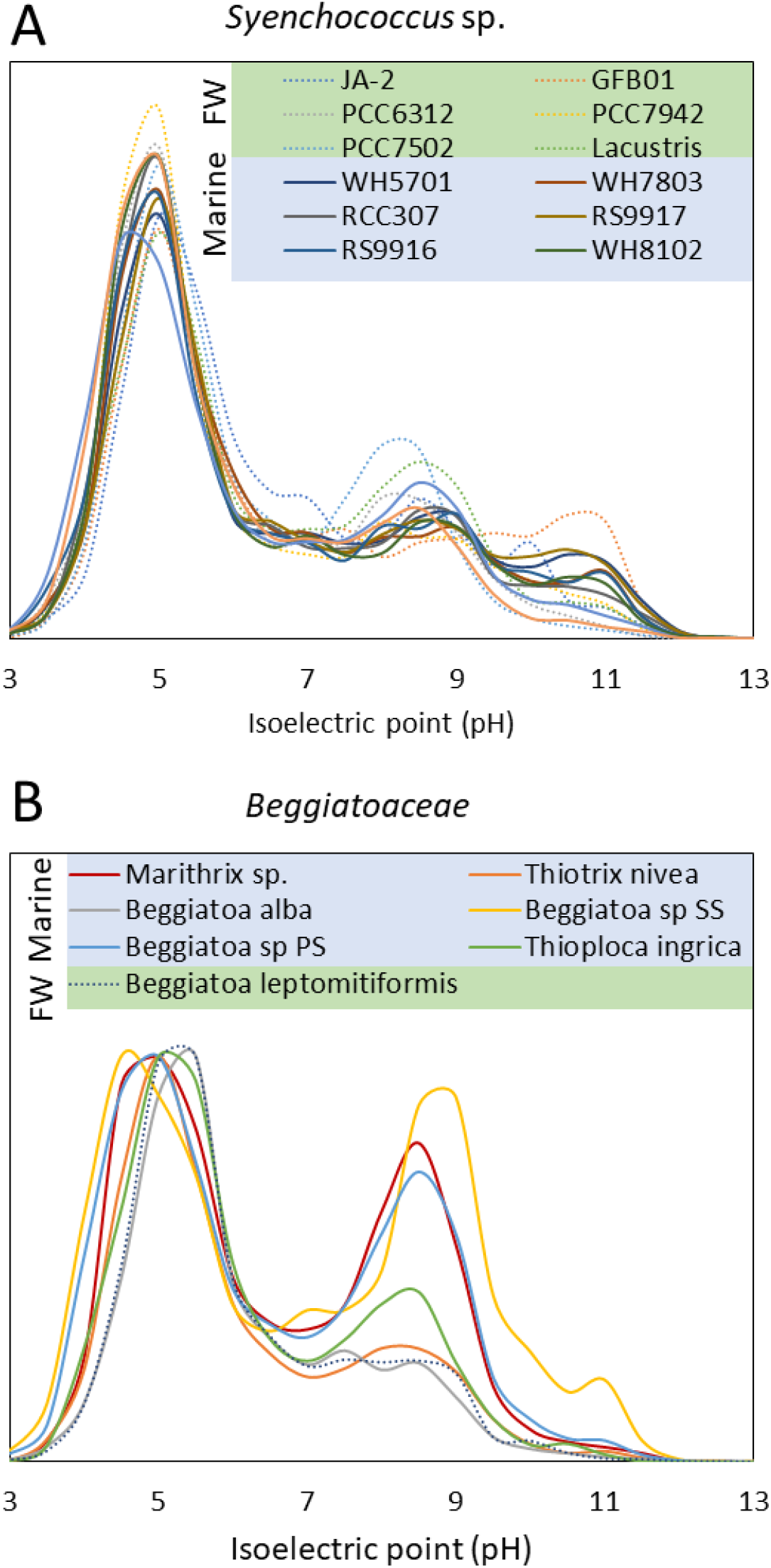
Isoelectric point histograms of freshwater (dashed lines) and marine (full lines) Synechococcus proteomes (A) and members of the *Beggiaoaceae*. The proteomes for *Synechococcus lacustris, Beggiatoa leptomitiformis* and *Marithrix* were obtained from Genbank and calculated using the standalone version of the isoelectric point calculator (Kozlowski 2016). The remaining datasets were obtained from the pre-calculated database Proteome-PI (Kozlowski 2017). *Beggiatoa* sp. SS was sampled from a brackish harbor in the Baltic Sea with a salinity of ca. 4.5 PSU.

**Figure S3.**
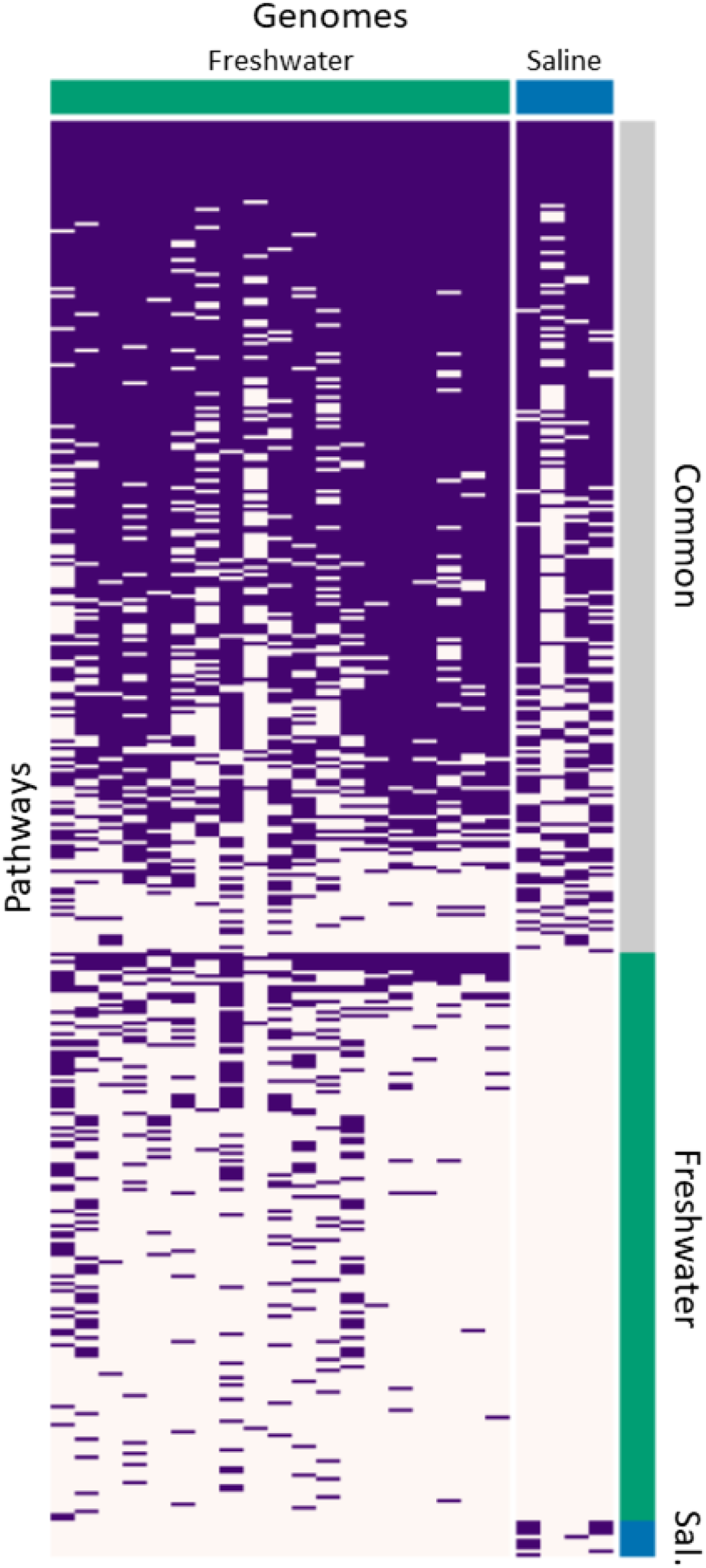
A comparison between functions detected in genomes from freshwater and marine (saline) *Achromatia*. The functions were predicted using Pathway Tools (Karp et al. 2015; Karp et al. 2019).

**Figure S4.**
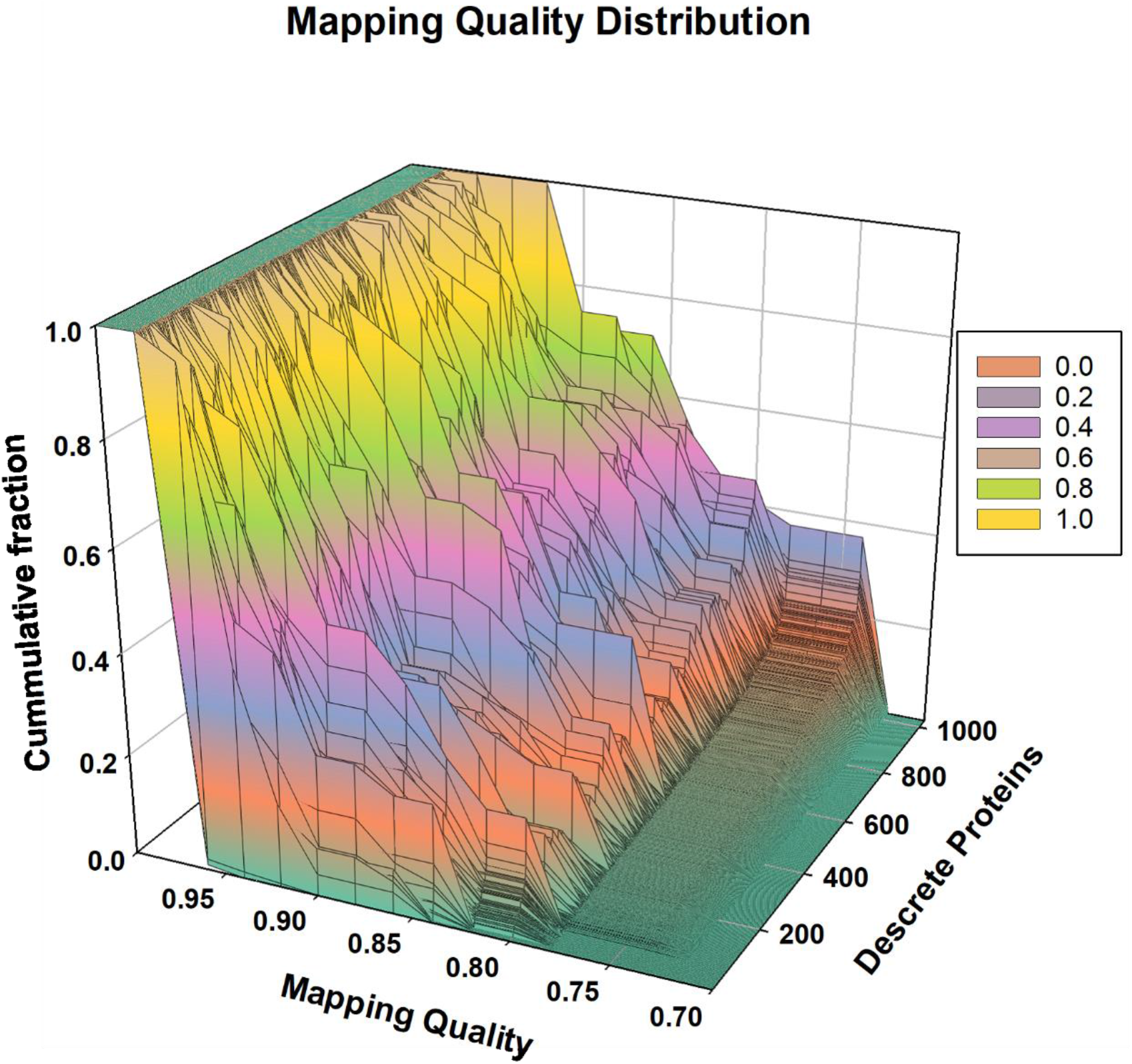
A set of histograms displaying the distribution of probabilities for a sequence from a raw dataset to correctly match an assembled gene from the *Achromatia* pangenome. Only genes (predicted proteins) occurring in more than 30 metagenomic datasets are shown. Probabilities (0.7-1) are displayed only for matches consider of sufficient quality to be displayed in Fig. 4 (i.e.>0.7).

**Figure S5.**
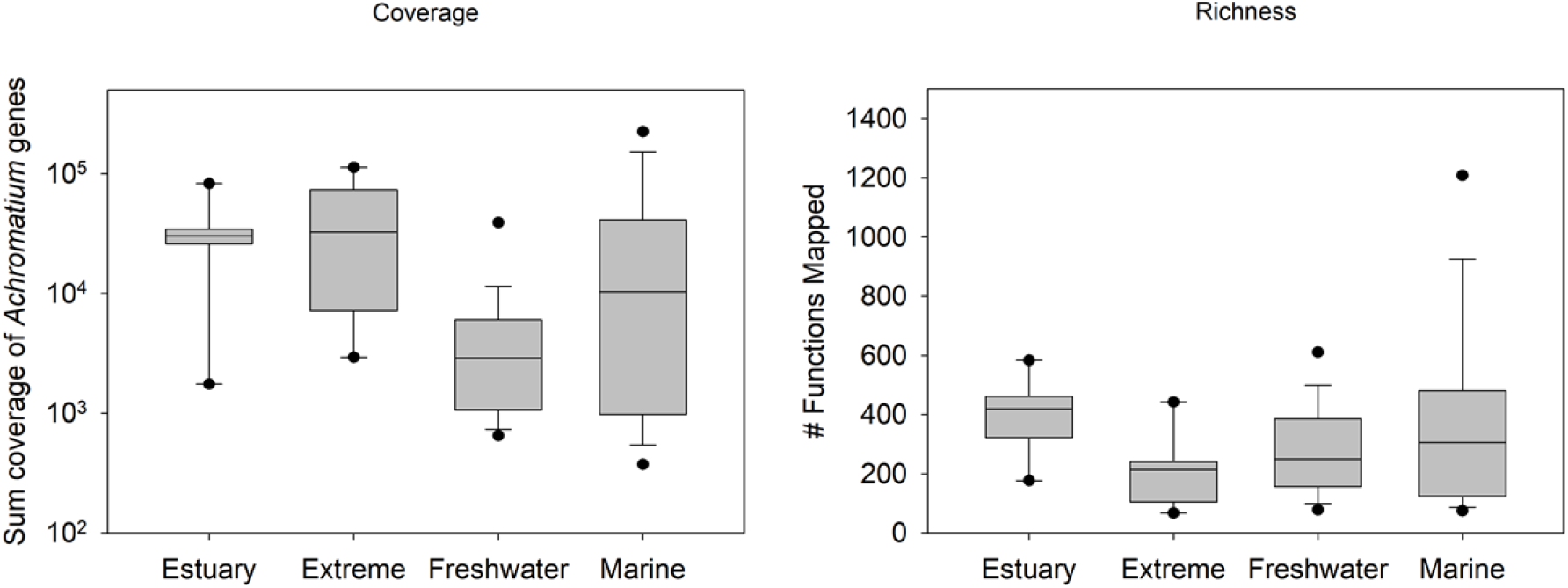
Distribution of functional coverage and richness values as calculated from mapping raw reads from public metagenomes to all predicted *Achromatia* genes.

**Figure S6.**
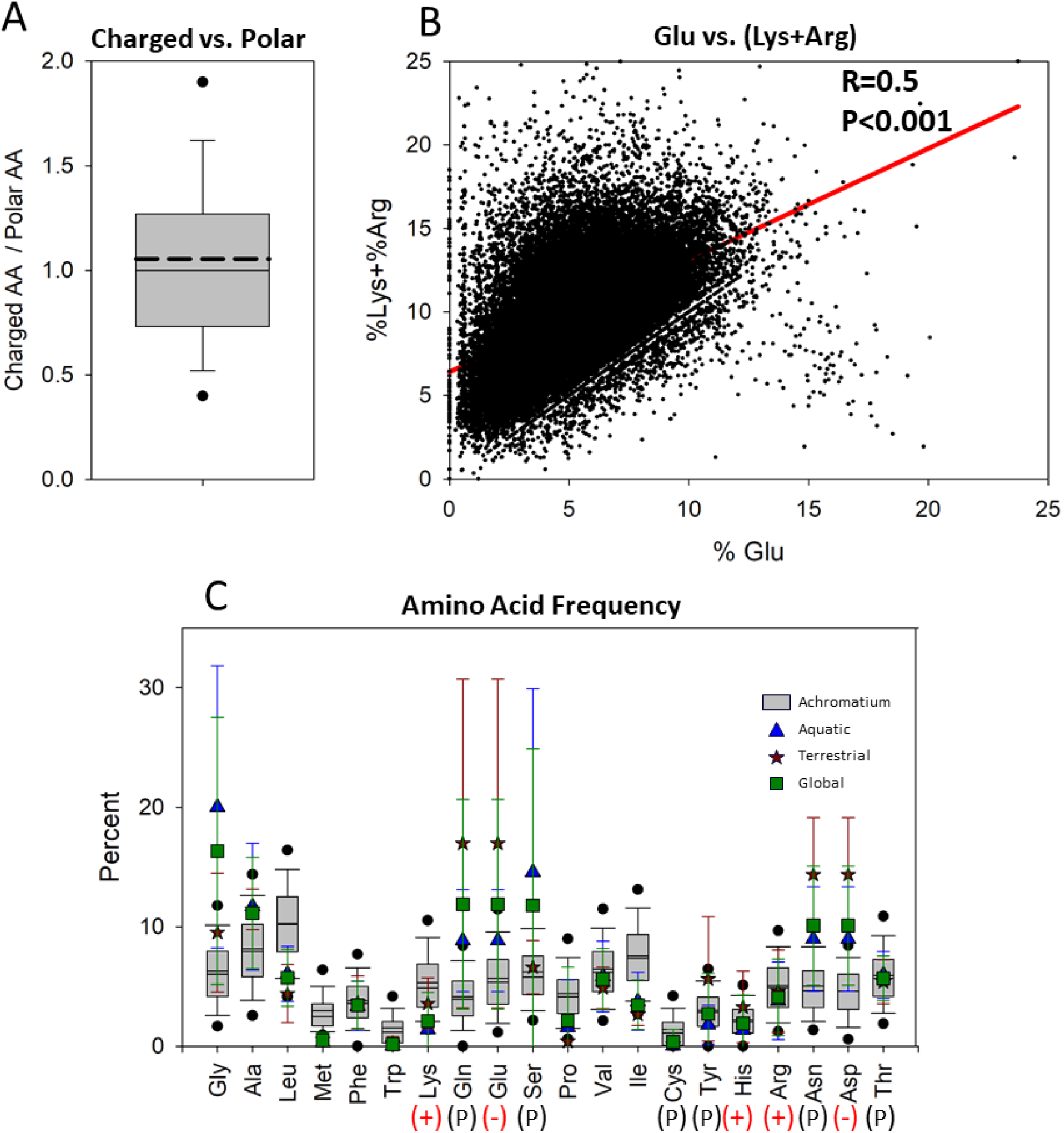
Analysis of Amino Acid frequency and patterns in the *Achromatia* pangenome. The ratio between charged and polar amino acids (A) is low typical of mesophilic bacteria with that of thermophiles being above 15. In contrast there is a thermophile-typical significant correlation between the frequency of Glu and the summed frequency of Lys and Arg (B). However, overall amino acid frequencies of the *Achromatia* genomes, differs from that of known aquatic and terrestrial bacteria (C) with higher frequencies of Leu, Lys and Ile and lower frequencies of Gln, Glu, Ser, Asn and Asp.

**Figure S7.**
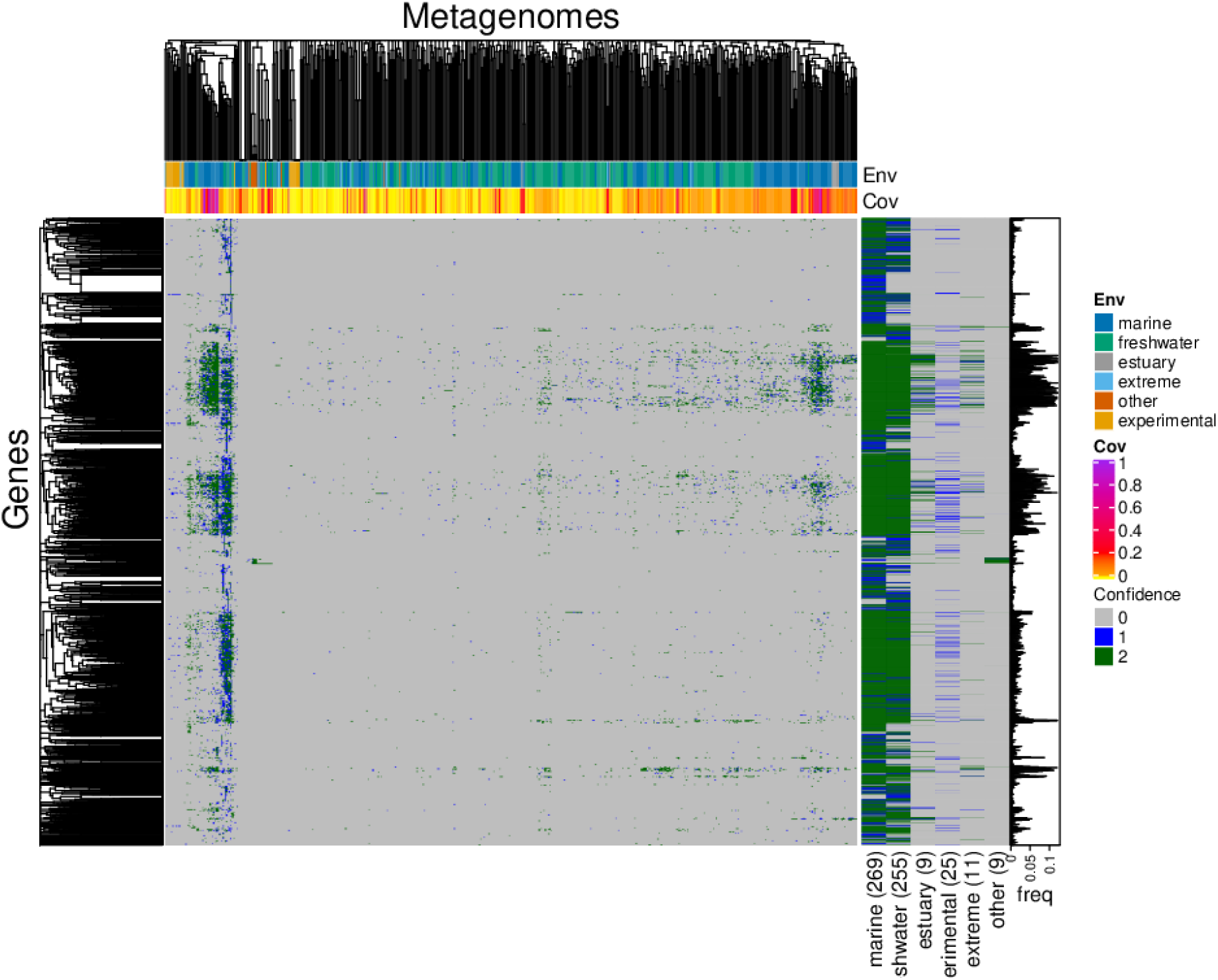
Continuation of Fig. 4, depicting functions with low abundance. The figures combined show the functional potential of *Achromatia* from different sediment environments as analyzed by mapping of raw sequence data to all known *Achromatia* annotated genes. Green color shows a probability of 75 % or higher that the mapping to a known *Achromatia* sequence is correct. Blue shows a lower probability while gray shows the function was not detected in the sample. Given that average amino acid identity between *Achromatia* from the same and different environments is as low as 65 % and 55 %, respectively (Ionescu et al. 2017), a match probability of 70 % and higher must be considered a high confidence match. The per-sample sum of the fold-coverage for each known *Achromatia* protein as calculated given by the BBMAP mapping program was used as a proxy for sequence coverage of *Achromatia* in the sample.

**Figure S8.**
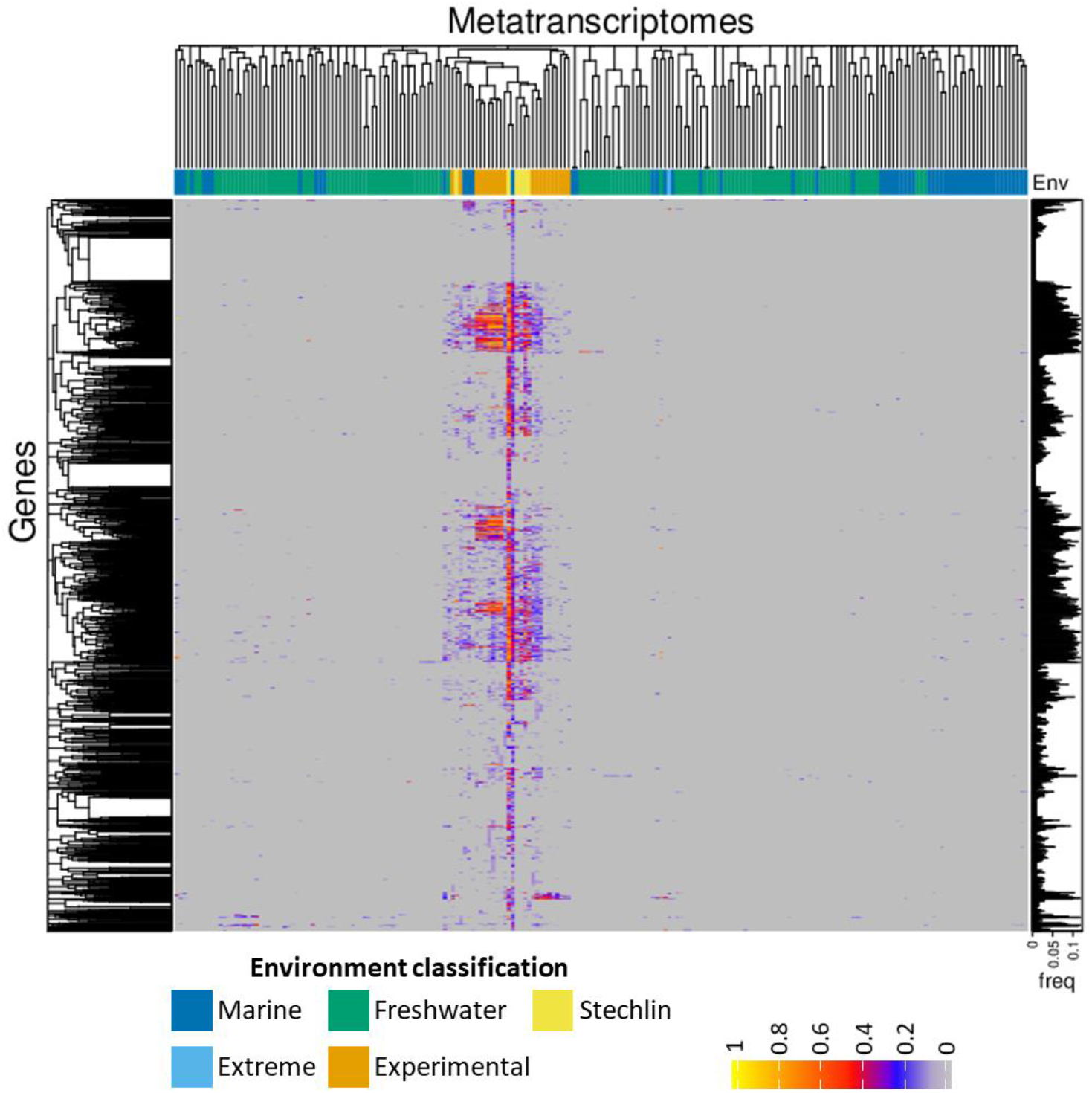
– Continuation of Fig. 5 showing rarer functions detected in the analysis of 300 publicly available sediment metatranscriptomes mapped to all known *Achromatia* functional genes and excluding ribosomal RNA and ribosomal proteins. To account for different sequencing depth and *Achromatia* cell abundance per sample, data was Log transformed and normalized per sample to range between 0-1.

**Figure S9.**
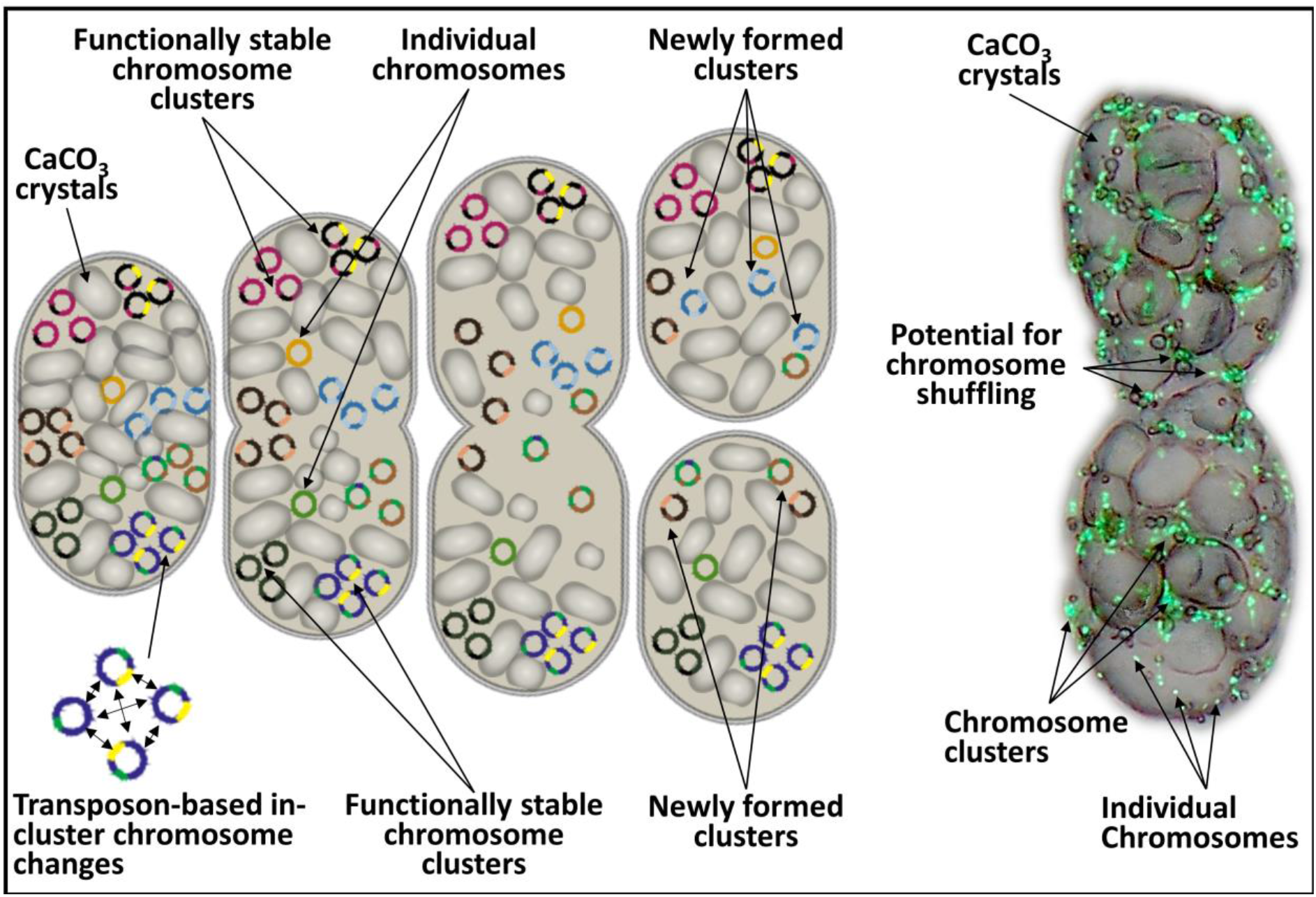
Hypothetical model of genetic diversity propagation in *Achromatia*, including the presence of genomic clusters alongside individual chromosomes. Transposon-based changes occur within each cluster. During cell division, some clusters are unaffected, providing the daughter cells with functional stability, while other clusters are shuffled leading to the formation of new clusters and thus eventually new chromosome versions. The overlaid light micrograph and DNA staining depicts chromosome clusters, individual chromosomes and sites of potential shuffling between clusters.

## References

Ahlgren NA, Rocap G. 2012. Diversity and Distribution of Marine Synechococcus: Multiple Gene Phylogenies for Consensus Classification and Development of qPCR Assays for Sensitive Measurement of Clades in the Ocean. Front. Microbiol. 3:213.

Aoshima M, Igarashi Y. 2006. A novel oxalosuccinate-forming enzyme involved in the reductive carboxylation of 2-oxoglutarate in Hydrogenobacter thermophilus TK-6. Mol. Microbiol. 62:748–759.

Babenzien H-D, Glöckner FO, Head IM. 2015. Achromatium. In: Bergey’s Manual of Systematics of Archaea and Bacteria. Chichester, UK: John Wiley & Sons, Ltd. p. 1–8.

Bankevich A, Nurk S, Antipov D, Gurevich AA, Dvorkin M, Kulikov AS, Lesin VM, Nikolenko SI, Pham S, Prjibelski AD, et al. 2012. SPAdes: a new genome assembly algorithm and its applications to single-cell sequencing. J. Comput. Biol. 19:455–477.

Batut B, Knibbe C, Marais G, Daubin V. 2014. Reductive genome evolution at both ends of the bacterial population size spectrum. Nat. Rev. Microbiol. 12:841–850.

Becraft ED, Woyke T, Jarett J, Ivanova N, Godoy-Vitorino F, Poulton N, Brown JM, Brown J, Lau MCY, Onstott T, et al. 2017. Rokubacteria: Genomic Giants among the Uncultured Bacterial Phyla. Front. Microbiol. 8:2264.

Bizic-Ionescu M, Ionescu D. 2016. Crossing the border - The freshwater/salt saline barrier: A phylogenetic analysis of bacteria inhabiting both freshwater and marine ecosystems. In: Glibert PM, Kanna T, editors. Aquatic Nutrient Biogeochemistry and Microbial Ecology: A Dual Perspective. Springer. p. in press.

Bleuven C, Landry CR. 2016. Molecular and cellular bases of adaptation to a changing environment in microorganisms. Proc. R. Soc. B Biol. Sci. 283.

Bobay L-M, Ochman H. 2017. The Evolution of Bacterial Genome Architecture. Front. Genet. 8:72.

Boscaro V, Felletti M, Vannini C, Ackerman MS, Chain PSG, Malfatti S, Vergez LM, Shin M, Doak TG, Lynch M, et al. 2013. Polynucleobacter necessarius, a model for genome reduction in both free-living and symbiotic bacteria. Proc. Natl. Acad. Sci. U. S. A. 110:18590–18595.

Cabello-Yeves PJ, Rodriguez-Valera F. 2019. Marine-freshwater prokaryotic transitions require extensive changes in the predicted proteome. Microbiome 7:117.

Cabello-Yeves PJ, Zemskay TI, Rosselli R, Coutinho FH, Zakharenko AS, Blinov V V., Rodriguez-Valera F. 2018. Genomes of novel microbial lineages assembled from the sub-ice waters of Lake Baikal. Appl. Environ. Microbiol. 84.

Cantarel BL, Coutinho PM, Rancurel C, Bernard T, Lombard V, Henrissat B. 2009. The Carbohydrate-Active EnZymes database (CAZy): an expert resource for Glycogenomics. Nucleic Acids Res. 37:D233–D238.

Christie-Oleza JA, Fernandez B, Nogales B, Bosch R, Armengaud J. 2012. Proteomic insights into the lifestyle of an environmentally relevant marine bacterium. ISME J. 6:124–135.

Dahl C, Engels S, Pott-Sperling AS, Schulte A, Sander J, Lübbe Y, Deuster O, Brune DC. 2005. Novel genes of the dsr gene cluster and evidence for close interaction of Dsr proteins during sulfur oxidation in the phototrophic sulfur bacterium Allochromatium vinosum. J. Bacteriol. 187:1392–1404.

Dvořák P, Casamatta DA, Poulíčková A, Hašler P, Ondřej V, Sanges R. 2014. *Synechococcus*: 3 billion years of global dominance. Mol. Ecol. 23:5538–5551.

Ghylin TW, Garcia SL, Moya F, Oyserman BO, Schwientek P, Forest KT, Mutschler J, Dwulit-Smith J, Chan LK, Martinez-Garcia M, et al. 2014. Comparative single-cell genomics reveals potential ecological niches for the freshwater acI Actinobacteria lineage. ISME J. 8:2503–2516.

Giovannoni SJ, Tripp HJ, Givan S, Podar M, Vergin KL, Baptista D, Bibbs L, Eads J, Richardson TH, Noordewier M, et al. 2005. Genome streamlining in a cosmopolitan oceanic bacterium. Science (80-.). 309:1242–1245.

Glöckner FO, Babenzien HD, Wulf J, Amann R. 1999. Phylogeny and diversity of Achromatium oxaliferum. Syst. Appl. Microbiol. 22:28–38.

Gray ND, Howarth R, Rowan A, Pickup RW, Jones JG, Head IM. 1999. Natural communities of Achromatium oxaliferum comprise genetically, morphologically, and ecologically distinct subpopulations. Appl. Environ. Microbiol. 65:5089–5099.

Grote J, Cameron Thrash J, Huggett MJ, Landry ZC, Carini P, Giovannoni SJ, Rappé MS. 2012. Streamlining and core genome conservation among highly divergent members of the SAR11 clade. MBio 3.

Gruber-Vodicka HR, Seah BK, Pruesse E. 2019. phyloFlash — Rapid SSU rRNA profiling and targeted assembly from metagenomes. bioRxiv:521922.

Gu Z, Eils R, Schlesner M. 2016. Complex heatmaps reveal patterns and correlations in multidimensional genomic data. Bioinformatics 32:2847–2849.

Guan N, Liu L. 2020. Microbial response to acid stress: mechanisms and applications. Appl. Microbiol. Biotechnol. 104:51–65.

Hartmann M, Howes CG, Abarenkov K, Mohn WW, Nilsson RH. 2010. V-Xtractor: An open-source, high-throughput software tool to identify and extract hypervariable regions of small subunit (16S/18S) ribosomal RNA gene sequences. J. Microbiol. Methods 83:250–253.

Häusler S, Weber M, de Beer D, Ionescu D. 2014. Spatial distribution of diatom and cyanobacterial mats in the Dead Sea is determined by response to rapid salinity fluctuations. Extremophiles 18:1085–1094.

Head IM, Gray ND, Howarth R, Pickup RW, Clarke KJ, Jones JG. 2000. Achromatium oxaliferum: Understanding the unmistakable. Adv. Microb. Ecol. 16:1–40.

Hellweger FL, Huang Y, Luo H. 2018. Carbon limitation drives GC content evolution of a marine bacterium in an individual-based genome-scale model. ISME J. 12:1180–1187.

Hong W, Wu YE, Fu X, Chang Z. 2012. Chaperone-dependent mechanisms for acid resistance in enteric bacteria. Trends Microbiol. 20:328–335.

Hottes AK, Freddolino PL, Khare A, Donnell ZN, Liu JC, Tavazoie S. 2013. Bacterial Adaptation through Loss of Function. PLoS Genet. 9.

Hurst LD, Merchant AR. 2001. High guanine–cytosine content is not an adaptation to high temperature: a comparative analysis amongst prokaryotes. Proc. R. Soc. London. Ser. B Biol. Sci. 268:493–497.

Ionescu D, Bizic-Ionescu M, De Maio N, Cypionka H, Grossart H-P. 2017. Community-like genome in single cells of the sulfur bacterium Achromatium oxaliferum. Nat. Commun. 8:455.

Ionescu D, Bizic M. 2019. Giant Bacteria. In: eLS. Chichester, UK: John Wiley & Sons, Ltd. p. 1–10.

Kanehisa M, Sato Y, Morishima K. 2016. BlastKOALA and GhostKOALA: KEGG Tools for Functional Characterization of Genome and Metagenome Sequences. J. Mol. Biol. 428:726–731.

Karp PD, Billington R, Caspi R, Fulcher CA, Latendresse M, Kothari A, Keseler IM, Krummenacker M, Midford PE, Ong Q, et al. 2019. The BioCyc collection of microbial genomes and metabolic pathways. Brief. Bioinform. 20:1085–1093.

Karp PD, Paley SM, Midford PE, Krummenacker M, Billington R, Kothari A, Ong WK, Subhraveti P, Keseler IM, Caspi R. 2015. Pathway Tools version 23.0: Integrated Software for Pathway/Genome Informatics and Systems Biology. Brief. Bioinform. 11:40–79.

Kashtan N, Roggensack SE, Rodrigue S, Thompson JW, Biller SJ, Coe A, Ding H, Marttinen P, Malmstrom RR, Stocker R, et al. 2014. Single-cell genomics reveals hundreds of coexisting subpopulations in wild Prochlorococcus. Science (80-.). 344:416–420.

Koonin E V. 2009. Evolution of genome architecture. Int. J. Biochem. Cell Biol. 41:298–306.

Kozlowski LP. 2016. IPC-Isoelectric Point Calculator. Biol. Direct 11.

Kozlowski LP. 2017. Proteome-pI: proteome isoelectric point database. Nucleic Acids Res. 45:D1112–D1116.

Kumar S, Nussinov R. 2001. How do thermophilic proteins deal with heat? Cell. Mol. Life Sci. 58:1216–1233.

Lagkouvardos I, Joseph D, Kapfhammer M, Giritli S, Horn M, Haller D, Clavel T. 2016. IMNGS: A comprehensive open resource of processed 16S rRNA microbial profiles for ecology and diversity studies. Sci. Rep. 6:1–9.

Ludt K, Soppa J. 2019. Polyploidy in halophilic archaea: regulation, evolutionary advantages, and gene conversion. Biochem. Soc. Trans. 47:933–944.

Lykidis D, Van Noorden S, Armstrong A, Spencer-Dene B, Li J, Zhuang Z, Stamp GWH. 2007. Novel zinc-based fixative for high quality DNA, RNA and protein analysis. Nucleic Acids Res. 35:e85–e85.

Lynch M. 2006. Streamlining and Simplification of Microbial Genome Architecture. Annu. Rev. Microbiol. 60:327–349.

Mansor M, Hamilton TL, Fantle MS, Macalady JL. 2015. Metabolic diversity and ecological niches of Achromatium populations revealed with single-cell genomic sequencing. Front. Microbiol. 6:822.

Markov A V, Kaznacheev IS. 2016. Evolutionary consequences of polyploidy in prokaryotes and the origin of mitosis and meiosis. Biol. Direct 11:28.

Mende DR, Bryant JA, Aylward FO, Eppley JM, Nielsen T, Karl DM, Delong EF. 2017. Environmental drivers of a microbial genomic transition zone in the ocean’s interior. Nat. Microbiol. 2:1367–1373.

Mendell JE, Clements KD, Choat JH, Angert ER. 2008. Extreme polyploidy in a large bacterium. Proc. Natl. Acad. Sci. 105:6730–6734.

Milner DS, Attah V, Cook E, Maguire F, Savory FR, Morrison M, Müller CA, Foster PG, Talbot NJ, Leonard G, et al. 2019. Environment-dependent fitness gains can be driven by horizontal gene transfer of transporter-encoding genes. Proc. Natl. Acad. Sci. U. S. A. 116:5613–5622.

Mirete S, Morgante V, González-Pastor JE. 2017. Acidophiles: Diversity and Mechanisms of Adaptation to Acidic Environments. In: Adaption of Microbial Life to Environmental Extremes. Cham: Springer International Publishing. p. 227–251.

Mock T, Otillar RP, Strauss J, McMullan M, Paajanen P, Schmutz J, Salamov A, Sanges R, Toseland A, Ward BJ, et al. 2017. Evolutionary genomics of the cold-adapted diatom Fragilariopsis cylindrus. Nature 541:536–540.

Mußmann M, Hu FZ, Richter M, de Beer D, Preisler A, Jørgensen BB, Huntemann M, Glöckner FO, Amann R, Koopman WJH, et al. 2007. Insights into the Genome of Large Sulfur Bacteria Revealed by Analysis of Single Filaments. Moran NA, editor. PLoS Biol. 5:e230.

Nercessian O, Noyes E, Kalyuzhnaya MG, Lidstrom ME, Chistoserdova L. 2005. Bacterial populations active in metabolism of C1 compounds in the sediment of Lake Washington, a freshwater lake. Appl. Environ. Microbiol. 71:6885–6899.

Newton RJ, Griffin LE, Bowles KM, Meile C, Gifford S, Givens CE, Howard EC, King E, Oakley CA, Reisch CR, et al. 2010. Genome characteristics of a generalist marine bacterial lineage. ISME J. 4:784–798.

Oliverio AM, Katz LA. 2014. The dynamic nature of genomes across the tree of life. Genome Biol. Evol. 6:482–488.

Oren A. 2013. Life at high salt concentrations, intracellular KCl concentrations, and acidic proteomes. Front. Microbiol. 4:315.

Pattaragulwanit K, Brune DC, Trüper HG, Dahl C. 1998. Molecular genetic evidence for extracytoplasmic localization of sulfur globules in Chromatium vinosum. Arch. Microbiol. 169:434–444.

Van de Peer Y, Mizrachi E, Marchal K. 2017. The evolutionary significance of polyploidy. Nat. Rev. Genet. 18:411–424.

Price MN, Dehal PS, Arkin AP. 2010. FastTree 2-Approximately maximum-likelihood trees for large alignments. PLoS One 5:e9490.

Richter K, Haslbeck M, Buchner J. 2010. The Heat Shock Response: Life on the Verge of Death. Mol. Cell 40:253–266.

Rodriguez-R L, Konstantinidis K. 2016. The enveomics collection: a toolbox for specialized analyses of microbial genomes and metagenomes.

Saarinen K, Laakso J, Lindström L, Ketola T. 2018. Adaptation to fluctuations in temperature by nine species of bacteria. Ecol. Evol. 8:2901–2910.

Salcher MM, Schaefle D, Kaspar M, Neuenschwander SM, Ghai R. 2019. Evolution in action: habitat transition from sediment to the pelagial leads to genome streamlining in Methylophilaceae. ISME J. 13:2764–2777.

Salman-Carvalho V, Fadeev E, Joye SB, Teske A. 2016. How Clonal Is Clonal? Genome Plasticity across Multicellular Segments of a “Candidatus Marithrix sp.” Filament from Sulfidic, Briny Seafloor Sediments in the Gulf of Mexico. Front. Microbiol. 7:1173.

Salman V, Amann R, Girnth A-C, Polerecky L, Bailey J V., Høgslund S, Jessen G, Pantoja S, Schulz-Vogt HN. 2011. A single-cell sequencing approach to the classification of large, vacuolated sulfur bacteria. Syst. Appl. Microbiol. 34:243–259.

Salman V, Berben T, Bowers RM, Woyke T, Teske A, Angert ER. 2016. Insights into the single cell draft genome of “Candidatus Achromatium palustre.” Stand. Genomic Sci. 11.

Salman V, Yang T, Berben T, Klein F, Angert E, Teske A. 2015. Calcite-accumulating large sulfur bacteria of the genus Achromatium in Sippewissett Salt Marsh. ISME J. 9:2503–2514.

Schorn S, Salman-Carvalho V, Littmann S, Ionescu D, Grossart H-P, Cypionka H. 2020. Cell Architecture of the Giant Sulfur Bacterium Achromatium oxaliferum: Extra-cytoplasmic Localization of Calcium Carbonate Bodies. FEMS Microbiol. Ecol. 96.

Seemann T. 2014. Prokka: rapid prokaryotic genome annotation. Bioinformatics 30:2068–2069.

Serres MH, Kerr ARW, McCormack TJ, Riley M. 2009. Evolution by leaps: Gene duplication in bacteria. Biol. Direct 4:46.

Soppa J. 2011. Ploidy and gene conversion in archaea. In: Biochemical Society Transactions. Vol. 39. Biochem Soc Trans. p. 150–154.

Suhre K, Claverie JM. 2003. Genomic correlates of hyperthermostability, an update. J. Biol. Chem. 278:17198–17202.

Swan BK, Tupper B, Sczyrba A, Lauro FM, Martinez-Garcia M, Gonźalez JM, Luo H, Wright JJ, Landry ZC, Hanson NW, et al. 2013. Prevalent genome streamlining and latitudinal divergence of planktonic bacteria in the surface ocean. Proc. Natl. Acad. Sci. U. S. A. 110:11463–11468.

Tekaia F, Yeramian E, Dujon B. 2002. Amino acid composition of genomes, lifestyles of organisms, and evolutionary trends: A global picture with correspondence analysis. Gene 297:51–60.

Teske A, Salman V. 2014. The family beggiatoaceae. In: The Prokaryotes: Gammaproteobacteria. Vol. 9783642389. Berlin, Heidelberg: Springer Berlin Heidelberg. p. 93–134.

Tomatis PE, Fabiane SM, Simona F, Carloni P, Sutton BJ, Vila AJ. 2008. Adaptive protein evolution grants organismal fitness by improving catalysis and flexibility. Proc. Natl. Acad. Sci. U. S. A. 105:20605–20610.

Trent JD, Gabrielsen M, Jensen B, Neuhard J, Olsen J. 1994. Acquired thermotolerance and heat shock proteins in thermophiles from the three phylogenetic domains. J. Bacteriol. 176:6148–6152.

Tsementzi D, Rodriguez-R LM, Ruiz-Perez CA, Meziti A, Hatt JK, Konstantinidis KT. 2019. Ecogenomic characterization of widespread, closely-related SAR11 clades of the freshwater genus “Candidatus Fonsibacter” and proposal of Ca. Fonsibacter lacus sp. nov. Syst. Appl. Microbiol. 42:495–505.

Walsh DA, Lafontaine J, Grossart HP. 2013. On the eco-evolutionary relationships of fresh and salt water bacteria and the role of gene transfer in their adaptation. In: Gophna U, editor. Lateral Gene Transfer in Evolution. Vol. ed Gophna. New York, NY: Springer. p. 55–77.

Wang R, Lin Jian Qiang, Liu XM, Pang X, Zhang CJ, Yang CL, Gao XY, Lin CM, Li YQ, Li Y, et al. 2019. Sulfur oxidation in the acidophilic autotrophic Acidithiobacillus spp. Front. Microbiol. 10:3290.

Wickham H. 2016. ggplot2 Elegant Graphics for Data Analysise. New York: Springer New York

Wiedenbeck J, Cohan FM. 2011. Origins of bacterial diversity through horizontal genetic transfer and adaptation to new ecological niches. FEMS Microbiol. Rev. 35:957–976.

Winkel M, Salman-Carvalho V, Woyke T, Richter M, Schulz-Vogt HN, Flood BE, Bailey J V, Mußmann M. 2016. Single-cell Sequencing of Thiomargarita Reveals Genomic Flexibility for Adaptation to Dynamic Redox Conditions. Front. Microbiol. 7:964.

Yang T, Teske A, Ambrose W, Salman-Carvalho V, Bagnell R, Nielsen LP. 2019. Intracellular calcite and sulfur dynamics of Achromatium cells observed in a lab-based enrichment and aerobic incubation experiment. Antonie Van Leeuwenhoek 112:263–274.

Zaremba-Niedzwiedzka K, Viklund J, Zhao W, Ast J, Sczyrba A, Woyke T, McMahon K, Bertilsson S, Stepanauskas R, Andersson SGE. 2013. Single-cell genomics reveal low recombination frequencies in freshwater bacteria of the SAR11 clade. Genome Biol. 14:R130.

Zhang H, Yohe T, Huang L, Entwistle S, Wu P, Yang Z, Busk PK, Xu Y, Yin Y. 2018. dbCAN2: a meta server for automated carbohydrate-active enzyme annotation. Nucleic Acids Res. 46:W95–W101.

